# Polymer-lipid hybrid nanoparticle enhances mRNA delivery and T cell-mediated immunity

**DOI:** 10.64898/2026.01.22.701138

**Authors:** Xiaolei Cai, Min Chen, Guoshuai Cao, Nicholas Asby, Derek Elli, Haley Gula, Vlad Nicolaescu, Duy-Thuc Nguyen, Xiaodan Huang, Tanushree Dangi, Ani Solanki, Sara Woessner, Wenbo Zhang, Erting Tang, Lisa Volpatti, Rachel Wallace, Tony Pan, Mindy Nguyen, Qing Chen, Zhi Geng, Rohin Sagar, Aaron Esser-Khan, Pablo Penaloza-MacMaster, Dominique Missiakas, Jun Huang

## Abstract

mRNA vaccines have transformed prophylactic immunization against infectious diseases as well as therapeutic interventions for cancer. However, their effectiveness against emerging viral variants and a range of malignancies continues to be hindered by suboptimal induction of T cell-mediated immunity. To overcome this limitation, here we developed a polymer-lipid hybrid nanoparticle (PLNP) platform engineered to improve mRNA delivery to antigen-presenting cells (APCs) and to potentiate T cell responses. Relative to conventional mRNA lipid nanoparticle (LNP) vaccines, mRNA PLNP vaccines demonstrated markedly improved lymph node targeting, APC activation, Th1-biased pro-inflammatory cytokine response, and antigen-specific T cell expansion while retaining robust humoral immunity. Remarkably, mRNA-PLNP vaccines generated approximately 50% more antigen-specific CD8^+^ T cells than mRNA-LNP vaccines across multiple antigens, including SARS-CoV-2 spike, influenza hemagglutinin, and ovalbumin. In prophylactic applications, mRNA PLNP vaccine provided complete protection against SARS-CoV-2 variants. As a therapeutic approach in a melanoma model, mRNA PLNP vaccination resulted in enhanced tumor control and significantly prolonged survival compared to LNP-based formulations. Collectively, these results establish PLNP as a versatile and broadly applicable platform for augmenting mRNA vaccine efficacy through improved mRNA delivery and T cell priming, offering promising implications for infection prevention and cancer immunotherapy.

## INTRODUCTION

The lipid nanoparticle (LNP)-based mRNA vaccine platform has profoundly transformed vaccine development.^1,2^ The LNP formulation is crucial for stabilizing mRNA, protecting it from enzymatic degradation, facilitating efficient cellular uptake, and promoting endosomal escape for translation within the cytoplasm.^3,4^ In addition to their role as delivery vehicles, LNPs function as self-adjuvants, actively stimulating innate immune pathways and enhancing pro-inflammatory cytokine secretion, which strengthens antigen presentation and helps initiate adaptive immune responses.^5,6^ Consequently, synthetic mRNA encoding specific antigens enables host cells to produce and present immunogenic proteins, driving strong adaptive immunity.^7,8^ The ability of LNPs to both enhance delivery and boost immune potency synergistically promotes antigen expression and immunogenicity, supports rapid antigen design and modification, and enables scalable manufacturing.^9,10^

The success of COVID-19 mRNA vaccines exemplifies the power of this technology, as these vaccines elicit strong humoral immunity, leading to the production of specific antibodies that bind and neutralize the virus, thereby preventing infection and protecting the host.^11–13^ However, despite these significant advances, current mRNA-LNP vaccines have limited capacity to induce robust cellular immunity.^14–17^ T cell-mediated immunity is critically important because, unlike antibodies that primarily block virus entry, T cells can directly recognize and eliminate infected cells, preventing viral spread.^18,19^ This cellular response is particularly important against emerging viral variants that may evade antibody recognition, given that T cell can recognize conserved linear epitopes present among many viral variants.^20–22^ Additionally, the activation of antigen-specific T cells is fundamental for long-term immune memory and for effective cancer immunotherapy, as these cells can target and destroy tumor cells.^23–25^ Therefore, efficient induction of T cell responses is essential not only for controlling evolving infections but also for achieving sustained tumor surveillance. There is an urgent need for next-generation mRNA delivery platforms that more effectively elicit T cell-mediated immunity, thereby enabling broader and more durable protection against both infection and cancer.

Efficient mRNA translation and antigen presentation in antigen-presenting cells (APCs) are crucial for priming and generating antigen-specific T cells.^26–28^ Building on our previous findings that certain polymers enhance nanoparticle uptake by APCs,^29,30^ we hypothesize that polymer-lipid hybrid nanoparticles (PLNPs) can provide better control over mRNA release, enhance APC uptake, and improve antigen presentation, while retaining essential advantages such as high mRNA encapsulation efficiency, low cytotoxicity, scalability in manufacturing, and broad compatibility with various mRNA constructs. Through a series of *in vitro* and *in vivo* development and evaluation of the PLNP delivery platform in both prophylactic (SARS-CoV-2 and influenza) and therapeutic (OVA-melanoma) settings, our results demonstrate that PLNPs are significantly superior to conventional LNPs, enabling more robust generation of antigen-specific T cells and markedly enhanced cell-mediated immune responses. Notably, PLNPs maintain a comparable level of humoral immune response to conventional LNPs, thereby preserving the benefits of strong antibody production. This approach offers broad potential applications across prophylactic vaccines, therapeutic cancer immunotherapies, *in vivo* cell therapies, and other mRNA-based therapies for different diseases.

## RESULTS

### Design principle and characterization of PLNP

Mounting an effective T cell-mediated immune response relies on mRNA delivery platforms that not only enhance mRNA delivery and translation for robust antigen presentation, but also efficiently activate APCs and produce pro-inflammatory cytokines necessary for Th1 cell polarization. Th1 cells are the key driver of cell-mediated immunity, acting primarily through the secretion of interferon-γ (IFN-γ) and activation of macrophages and cytotoxic CD8 T cells.^31,32^ Conventional LNPs, while highly successful in stabilizing and delivering mRNA, have limitations in inducing strong T cell-mediated immunity that is critical for lasting protection and cancer immunotherapy. Previous studies, including our own, have demonstrated that polymer-based nanoparticles significantly improve APC uptake, antigen delivery, and Th1-biased immune responses.^29,30,33–35^ Building on these insights, we hypothesize that engineering a PLNP platform can synergistically combine the strengths of both polymers and lipids to further enhance mRNA delivery, APC activation, and T cell priming, ultimately producing more potent and sustained antigen-specific T cell responses than conventional LNPs (**Scheme S1a**).

To test our hypothesis, we first designed a prototype polymer-lipid hybrid formulation (PLNP-0) using FDA-approved materials: an ionizable lipid (SM-102), a PEG-lipid (DMG-PEG), and a polymer, poly(lactic-co-glycolic) acid (PLGA) (**Scheme S1b**). The ionizable lipid enables efficient mRNA encapsulation and endosomal escape, supporting successful mRNA translation within the cell, and also stimulates innate immune pathways leading to the production of pro-inflammatory cytokines.^36,37^ The PEG-lipid helps to stabilize the nanoparticle, regulate its size, and prolong its circulation time.^3,38^ Most importantly, the addition of PLGA polymer is central to this technology, as it markedly increases nanoparticle stiffness, promoting cellular uptake by APCs and improving retention within draining lymph nodes (dLNs),^39–41^ which are the key sites for initiating adaptive immune responses. This approach should enhance mRNA delivery to dLNs, potentially boosting both the magnitude and longevity of T cell-mediated immunity beyond conventional LNPs.

After loading SARS-CoV-2 spike mRNA into both PLNP-0 and LNP formulations, we first assessed mRNA translation efficiency and immunostimulatory function in RAW 264.7 macrophages. As conserved T cell epitopes have been reported to confer broad protection through cell-mediated immunity,^42,43^ we included both ancestral SARS-CoV-2 (wild-type, WT) spike mRNA and B.1.1.529 spike mRNA (1:1) in the nanoparticle formulations to enhance coverage of the conserved T cell epitopes and thereby promote broader immunity against different variants. PLNP-0 demonstrated superior spike mRNA translation efficiency and higher secretion of cytokines, including IFN-γ, tumor necrosis factor-alpha (TNF-α), and interleukin-2 (IL-2), compared to LNPs (**Figs. S1a-d**). However, this promising *in vitro* performance did not translate to *in vivo* efficacy, as PLNP-0 failed to elicit anti-spike antibodies or spike-specific CD8^+^ T cell responses in mice (**Figs. S1e-f**). Investigation revealed that PLNP-0 has relatively low colloid stability, evidenced by altered nanoparticle size and increased polydispersity index after storage at 4 °C for 24 hours (**Figs. S1g-h**), which likely compromised its immune response *in vivo*. To address this, we improved the formulation by incorporating a helper lipid (1,2-distearoyl-sn-glycero-3-phosphocholine, DSPC) and cholesterol (**Scheme S1c**), yielding a more stable PLNP (**Figs. 1a, S2**). The refined PLNP exhibited a hydrodynamic diameter of approximately 150 nm, which is larger than LNPs (about 90 nm), indicating successful polymer incorporation (**Fig. 1b, left panel**). DLS measurements showed that both LNP and PLNP exhibited uniform size distribution, each with a polydispersity index (PDI) less than 0.2 (**Fig. 1c**), and transmission electron microscopy further confirmed their spherical morphology and comparable particle sizes (**Fig. 1b, right panel**). In terms of mRNA encapsulation, the efficiencies were similar (92% for LNP and 90% for PLNP, **Fig. 1d**).

**Figure 1.**
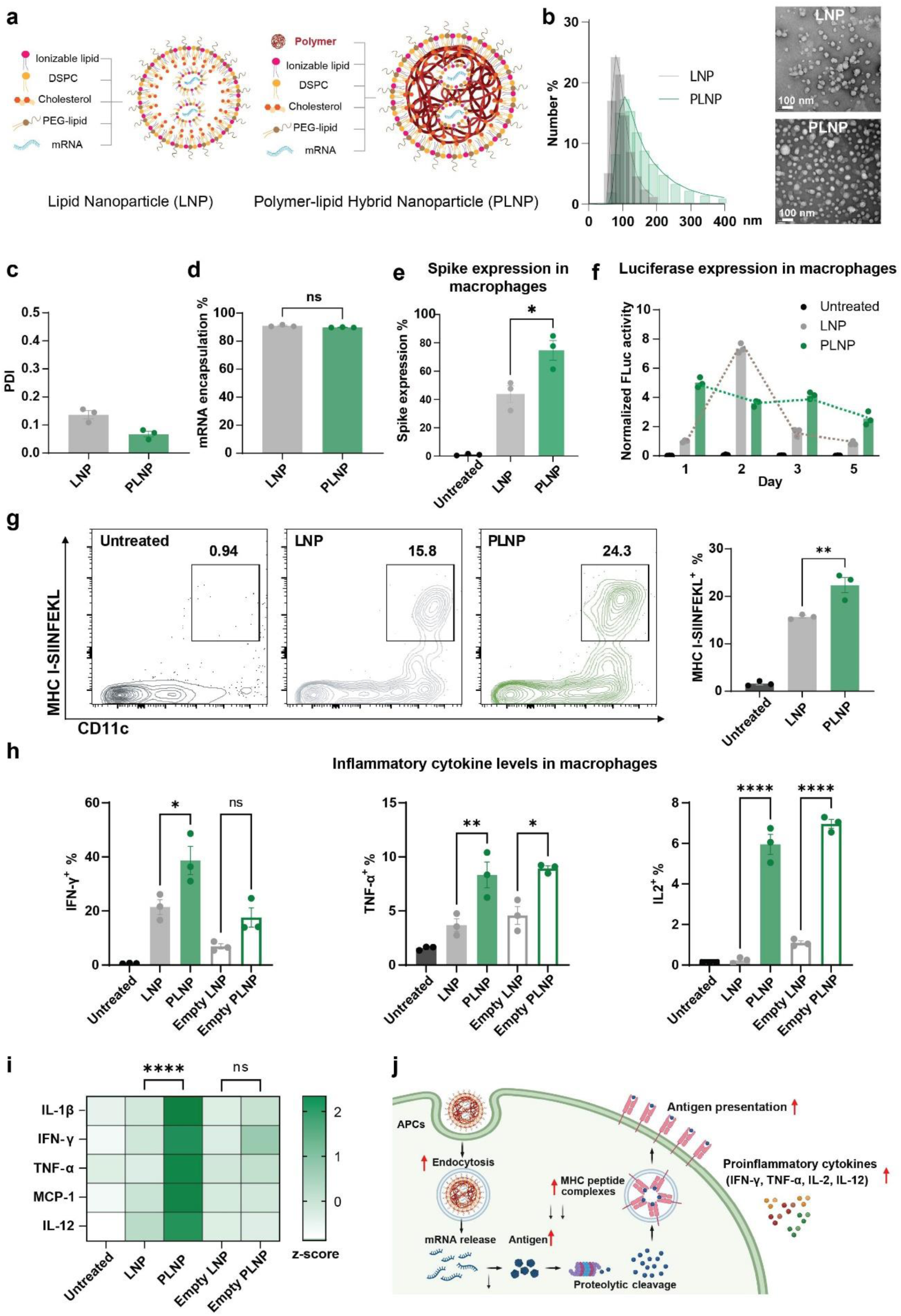
Design and *in vitro* characterization of PLNP. (a) Schematic illustration of the design of LNP and PLNP. (b) Left: sizes of LNP and PLNP measured by dynamic light scattering (DLS). Right: TEM images of LNP and PLNP. Scale bars: 100 nm. (c) Polydispersity index (PDI) of LNP and PLNP. (d) mRNA encapsulation efficiencies of LNP and PLNP. (e) Spike mRNA translation in RAW 264.7 macrophages after incubation with spike mRNA-loaded LNP or PLNP for 48 hours. Spike expression were measured by flow cytometry. Dose: 1 µg mRNA/mL. n = 3. (f) FLuc mRNA translation in RAW 264.7 macrophages on day 1, 2, 3, and 5 after incubation with FLuc mRNA-loaded LNP or PLNP. Dose: 1 µg mRNA/mL. n = 3. The FLuc signals were normalized to the day-1 signal of LNP-treated group. (g) Left: Representative flow plots of CD11c^+^ DCs expressing SIINFEKL/H-2K^b^ after incubation with OVA mRNA-loaded LNP or PLNP for 48 hours. Right: The percentages of CD11c^+^ SIINFEKL/H-2K^b+^ DCs in total DCs. Dose: 1 µg mRNA/mL. n = 3. (h) Intracellular cytokine levels in RAW 264.7 macrophages after incubation with spike mRNA-loaded LNP, spike mRNA-loaded PLNP, empty LNP, or empty PLNP for 24 hours. Left: IFN-γ. Middle: TNF-α. Right: IL-2. Dose: 1 µg mRNA/mL. n = 3. (i) Cytokine secretion from human PBMCs after incubation with spike mRNA-loaded LNP, spike mRNA-loaded PLNP, empty LNP, or empty PLNP for 24 hours. Dose: 1 µg mRNA/mL. n = 3. (j) Schematic illustration of PLNP enhancing antigen presentation and proinflammatory cytokine generation in APCs. Data are presented as mean ± s.e.m. Statistical significance was determined using two-sided unpaired t-test (d, e, g), one-way ANOVA with Tukey post hoc test for multiple comparisons (h), or two-way ANOVA with Tukey post hoc test for multiple comparisons ns: no significance, * P < 0.05, ** P < 0.01, *** P < 0.001, and ****P < 0.0001.

### PLNP enhances mRNA translation and antigen presentation *in vitro*

We first sought to determine whether PLNP could enhance mRNA translation efficiency and antigen presentation compared to conventional LNP, as antigen presentation is the first signal necessary for the activation of adaptive immune responses.^44^ RAW 264.7 macrophages were incubated with spike mRNA-loaded LNP or PLNP. While both formulations enabled cellular uptake and protein expression, PLNP-treated cells exhibited a significantly higher level of spike protein expression than those treated with LNP (**Fig. 1e**). This suggests that PLNP can facilitate better intracellular delivery and translation of mRNA cargo. Further analysis using firefly luciferase (FLuc) mRNA revealed that PLNP not only accelerated initial translation but also maintained stable and sustained expression over multiple days, whereas LNP-transfected cells showed a marked decline after day 2 **(Fig. 1f**). To further evaluate antigen presentation, bone marrow-derived dendritic cells (DCs) were exposed to OVA mRNA-loaded nanoparticles and assessed for peptide-major histocompatibility complex (pMHC) SIINFEKL/H-2K^b^ levels (**Fig. S3**). PLNP treatment resulted in a substantially greater frequency of SIINFEKL/H-2K^b^-positive DCs compared to LNP (**Fig. 1g**). These findings demonstrate that PLNP supports efficient and prolonged display of mRNA-encoded antigens, an essential prerequisite for effective T cell priming.

Since cytokine response is an important marker of APC activation and an essential signal for the activation of adaptive immune responses, we sought to determine whether PLNP could enhance immunostimulatory cytokine levels compared to LNP. PLNP-treated RAW 264.7 macrophages exhibited elevated levels of IFN-γ, TNF-α, and IL-2 at 24 hours post-treatment compared to LNP (**Fig. 1h**). The superiority of PLNP in stimulating APCs was confirmed in both human and mouse peripheral blood mononuclear cells (PBMCs), which produced higher levels of interleukin-1 beta (IL-1β), IFN-γ, and interleukin-12 (IL-12) in response to PLNP (**Figs. 1i, S4**), indicating potent induction of pro-inflammatory and Th1-skewed immune responses across species.

Collectively, these *in vitro* data demonstrate that PLNP is markedly more effective than conventional LNP in promoting mRNA translation, antigen presentation, and immunostimulatory cytokine production *in vitro* (**Fig. 1j**).

### PLNP promotes mRNA translation, lymph node targeting, and APC activation *in vivo*

To rigorously assess the potential of PLNP for mRNA delivery *in vivo*, we systematically compared its safety and functional performance against conventional LNP formulations. Importantly, all constituent lipids and polymer used for PLNP fabrication are FDA-approved, facilitating potential clinical translation. Given the larger size of PLNPs (**Fig. 1b**), we first evaluated biocompatibility by intramuscularly injecting C57BL/6 mice with either PLNP or LNP formulations and monitoring changes in body weight over a 28-day period. Both nanoparticle types exhibited negligible impact on body weight compared to the PBS control, confirming a favorable safety profile (**Fig. S5**).

We next assessed the efficiency of mRNA translation *in vivo* by immunizing BALB/c mice intramuscularly with FLuc mRNA-loaded nanoparticles and quantifying FLuc expression longitudinally. At four hours post-injection, PLNP-immunized mice exhibited nearly eight-fold higher bioluminescence, compared to those immunized with LNP, indicating enhanced early mRNA translation and antigen expression. Although signal intensity declined for both formulations at later time points, PLNP maintained significantly higher bioluminescence throughout the observation period, underscoring its superior capacity for promoting mRNA translation and generating more durable antigen expression *in vivo* (**Fig. 2a**).

**Figure 2.**
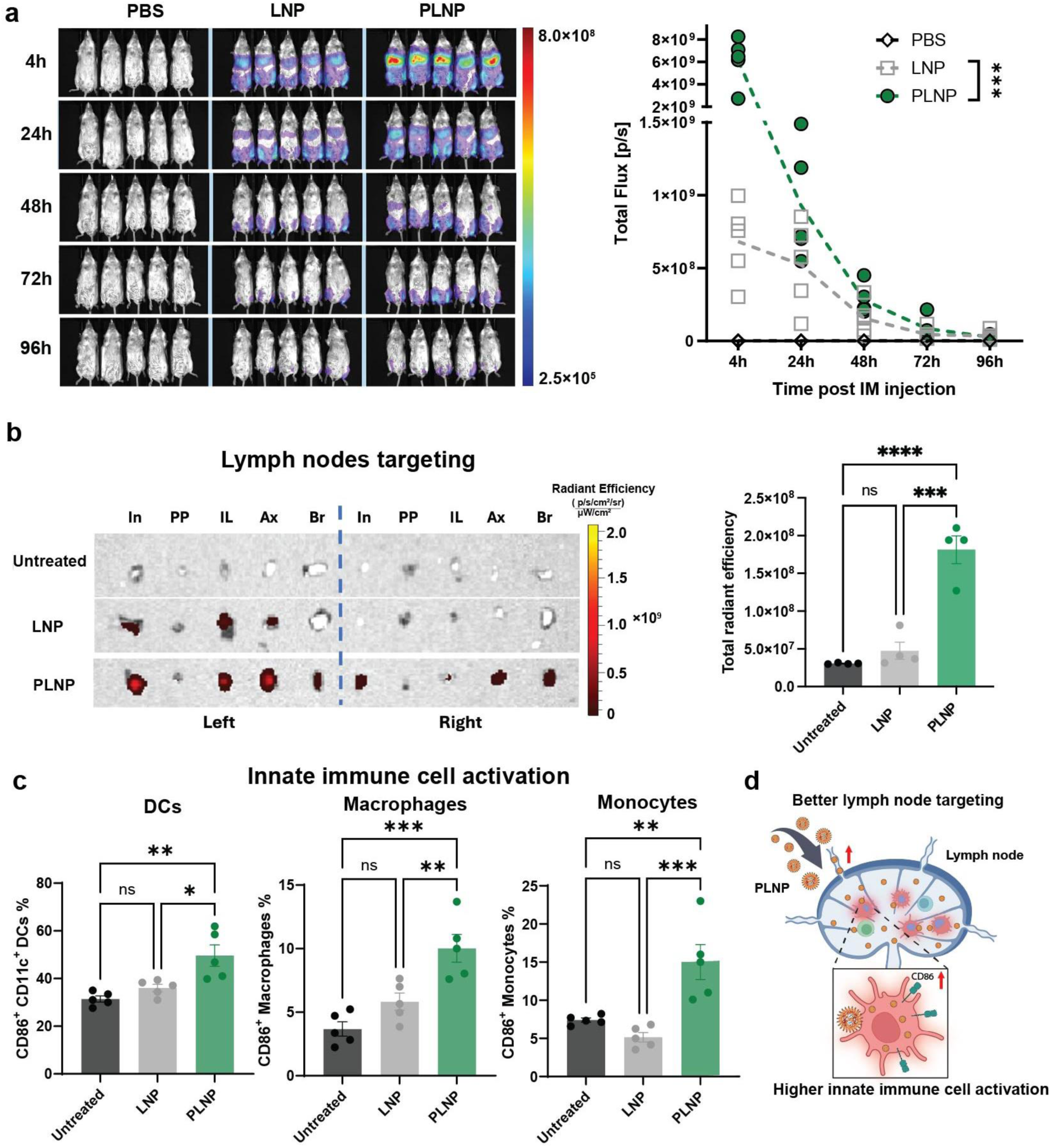
*In vivo* mRNA translation, lymph node targeting, and innate immune cell response after vaccination with LNP and PLNP. (a) Left: *In vivo* luminescence images of BALB/c mice treated with 1×PBS, FLuc mRNA-loaded LNP, or FLuc mRNA-loaded PLNP at 4, 24, 48, 72, and 96 hours post administration. Right: *In vivo* luminescence signals measured by integration of total flux for each mouse. Dose: 5 µg mRNA/mouse. N = 5. (b) Left: Fluorescence images of lymph nodes collected from C57BL/6 mice at 24 h post vaccination with DiD-labeled spike mRNA-loaded LNP or PLNP. Right: Quantification of LNP and PLNP accumulation in lymph nodes at 24 h post vaccination. Dose: 10 µg mRNA/mouse. n = 4. (c) Innate immune cell activation (CD86^+^) at 7 days post boost vaccination. C57BL/6 mice were vaccinated with spike mRNA-loaded LNP or PLNP on day 0 and boosted with same doses on day 21. Dose: 10 µg mRNA/mouse. n = 5. Left: DCs. Middle: Macrophages. Right: Monocytes. (d) Schematic illustration of PLNP enhancing lymph node targeting and innate immune cell activation. Data are presented as mean ± s.e.m. Statistical significance was determined using two-way ANOVA with Tukey post hoc test for multiple comparisons (a) or one-way ANOVA with Tukey post hoc test for multiple comparisons (b, c). ns: no significance, * P < 0.05, ** P < 0.01, *** P < 0.001, and ****P < 0.0001.

We then evaluated lymph node targeting, which is essential for robust immune activation and effective T cell priming; these tissues are the primary sites for antigen presentation and naïve T cell activation.^45^ Both PLNP and LNP were labeled with a fluorescent dye, 1,1’-dioctadecyl-3,3,3’,3’-tetramethylindodicarbocyanine, 4-chlorobenzenesulfonate salt (DiD), and administered intramuscularly to into the left quadriceps of C57BL/6 mice. At 24 hours post-injection, draining lymph nodes were harvested and analyzed for fluorescence to quantify nanoparticle accumulation. While both LNP and PLNP trafficked to regional lymph nodes, PLNP demonstrated broader distribution across a wider array of lymph nodes, including both ipsilateral and contralateral lymph nodes. Quantitative fluorescence analysis revealed that the signal intensities in lymph nodes were significantly higher in PLNP-treated mice compared to the LNP group, demonstrating that PLNP provides superior lymph node targeting (**Fig. 2b**).

Given that co-stimulation is another essential signal for the activation of adaptive immune responses, we sought to determine whether PLNP could enhance co-stimulation compared to LNP. We analyzed the expression of the co-stimulatory marker CD86 in DCs, macrophages, and monocytes following mRNA vaccination (**Fig. S6**). PLNP treatment led to stronger activation and higher expression of CD86 in these APC populations compared to LNP (**Fig. 2c**). In addition, PLNP-treated mice generally exhibited elevated systemic serum levels of pro-inflammatory cytokines, such as IL-23, IFN-γ, IL-12, and IFN-β, at 24 hours post-vaccination, reflecting more robust innate immune stimulation (**Fig. S7**). In addition, PLNP resulted in lower systemic IL-6 levels compared to LNP (**Fig. S7**), suggesting that PLNP could potentially reduce common systemic side effects associated with mRNA-LNP vaccines, such as fever and myalgia,^46^ and serve as a more tolerable formulation.

Collectively, these data demonstrate that PLNP is safe and markedly enhances mRNA translation, antigen expression, lymph node targeting, and APC co-stimulation compared with conventional LNP *in vivo* (**Fig. 2d**).

### PLNP triggers superior antigen-specific T cell immunity

We next investigated whether PLNP could elicit superior antigen-specific T cell immunity compared to LNP. mRNAs encoding SARS-CoV-2 spike (spike), influenza hemagglutinin (HA), and ovalbumin (OVA) proteins were individually encapsulated into LNP or PLNP formulations and administered intramuscularly to mice on day 0 and day 21. Mouse spleens were collected on day 28, and single-cell splenocyte suspensions were prepared to evaluate antigen-specific T cell responses.

To assess antigen-specific T cell immune responses, splenocytes were stimulated *ex vivo* with peptide pools corresponding to each antigen (spike, HA, or OVA), mixtures of overlapping peptides spanning the entire protein sequence, thereby ensuring broad activation of both CD4^+^ and CD8^+^ antigen-specific T cells. IFN-γ, TNF-α, and IL-2 secretions following stimulation were quantified using the LEGENDplex assay. Across all three antigens, splenocytes from PLNP-treated mice generally secreted higher levels of cytokines compared to those from LNP-treated mice, except for TNF-α in response to OVA. These results indicate more robust antigen-specific T cell responses in the PLNP-treated group (**Figs. 3a, c, e**).

**Figure 3.**
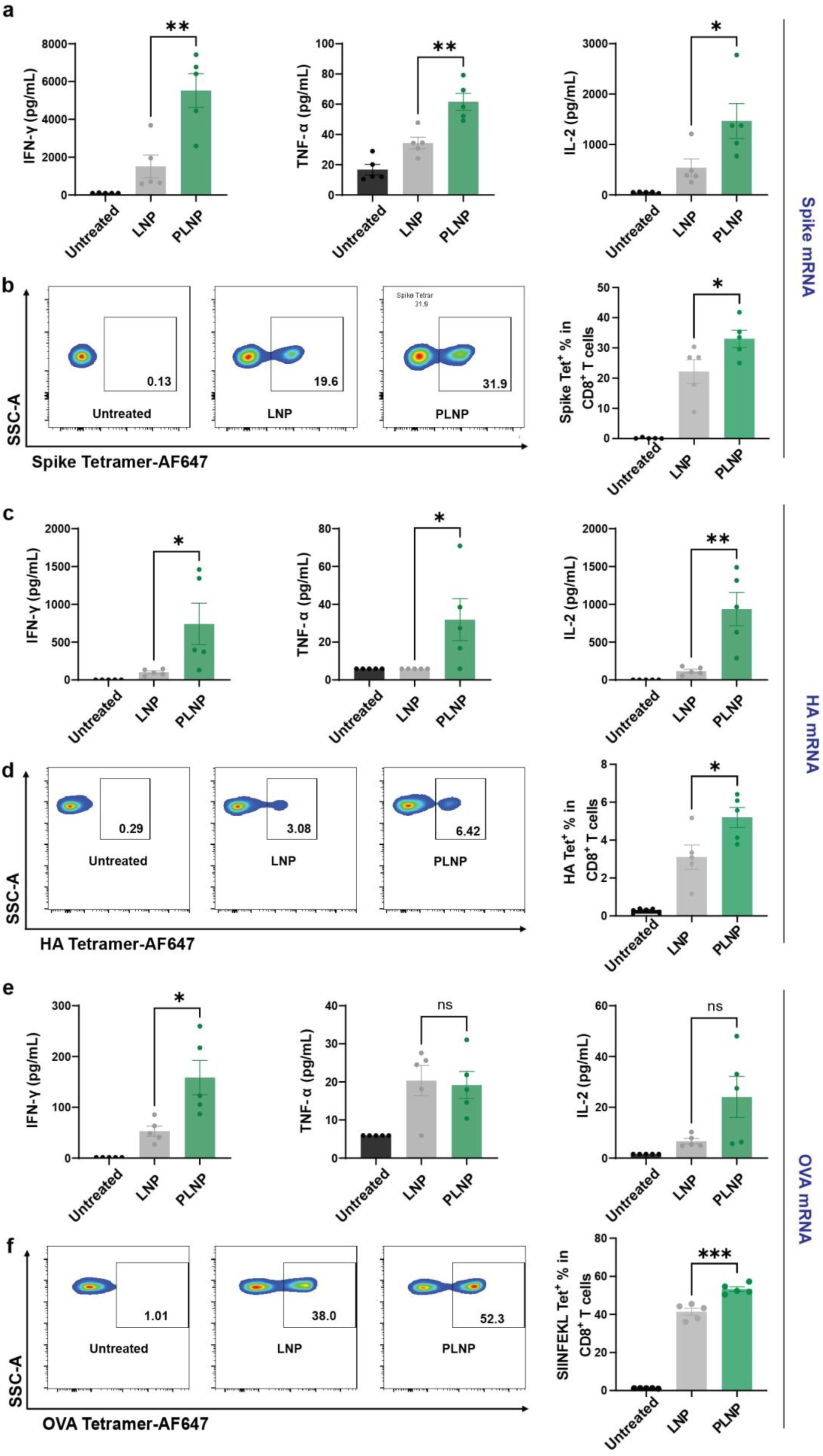
T-cell response post vaccination with LNP and PLNP. (a, c, e) IFN-γ, TNF-α, and IL-2 secretion after 6-hour peptide pools stimulation (spike, HA, or OVA) of the splenocytes collected on day 28. Mice were vaccinated with spike, HA, or OVA mRNA-loaded LNP or PLNP on day 0 and day 21. Dose: 10 µg spike mRNA, 10 µg HA mRNA, or 5 µg OVA mRNA/mouse. n = 5. (b, d, f) Left: Representative flow plots of antigen-specific CD8^+^ T cells in mouse splenocytes collected on day 28. Mice were vaccinated with spike, HA, or OVA mRNA-loaded LNP or PLNP on day 0 and day 21. Dose: 10 µg spike mRNA, 10 µg HA mRNA, or 5 µg OVA mRNA/mouse. n = 5. Antigen-specific CD8^+^ T cells are gated by tetramer^+^ cells in total CD8^+^ T cells. Right: The percentages of antigen-specific CD8^+^ T cells in total CD8^+^ T cells. Data are presented as mean ± s.e.m. Statistical significance was determined using two-sided unpaired t-test. ns: no significance, * P < 0.05, ** P < 0.01, and *** P < 0.001.

To further quantify antigen-specific CD8⁺ T cells, we performed pMHC tetramer staining.^47^ Splenocytes were incubated with antigen-specific pMHC tetramers (spike: VNFNFNGL/H-2K^b^, HA: IYSTVASSL/H-2K^d^, and OVA: SIINFEKL/H-2K^b^) and analyzed by flow cytometry to enumerate antigen-specific CD8⁺ T cells (**Fig. S8**). Consistent with the peptide pool stimulation results, PLNP treatment resulted in significantly greater numbers of antigen-specific CD8⁺ T cells across all three antigens (spike tetramer^+^: ∼33.0%, HA tetramer^+^: ∼5.2%, OVA tetramer^+^: ∼53.3%) compared to LNP (spike tetramer^+^: ∼22.2%, HA tetramer^+^: ∼3.1%, OVA tetramer^+^: ∼41.4%) (**Figs. 3b, d, f**). These results demonstrate that PLNP significantly enhances antigen-specific T cell immunity relative to conventional LNP, supporting its potential as an improved mRNA vaccine delivery platform.

### PLNP promotes CD8⁺ effector functions and Th1 polarization

In addition to specificity, we conducted a comprehensive analysis of T cell functional phenotypes following mRNA vaccination. C57BL/6 mice were immunized twice on days 0 and 21 with spike mRNA-loaded LNP or PLNP. Mouse spleens were harvested on day 28, and single-cell suspensions were prepared. Live CD8^+^ and CD4^+^ T cells were then single-cell sorted by fluorescence-activated cell sorting (FACS) and subjected to single-cell RNA sequencing using 10x Genomics kits.^48^

We first focused our analysis on CD8^+^ T cells. Following stringent quality control and filtering of the single-cell RNA sequencing data, dimensionality reduction was performed on the CD8^+^ T cell transcriptomes, and Louvain clustering was used to define major subpopulations. Eight distinct CD8^+^ T cell clusters were identified: naïve cells (Naïve), naïve cells with an early activation signature (Naïve_Act), central memory cells (CM), precursor effector cells (Eff_pre), effector cells expressing Il7r (Eff_Il7r), Runx2 (Eff_Runx2), and Btf3 (Eff_Btf3), as well as proliferative cells (Prolif) (**Figs. 4a, S9a**). Both mRNA-LNP and -PLNP vaccines induced CD8^+^ T cell repertoire changes compared to untreated group; however, PLNP triggered more significant changes (**Fig. 4a**, bottom panel). Most notably, PLNP led to a more than two-fold increase in the proportion of Runx2^+^ effector T cells (∼55%) relative to LNP (∼22%), underscoring the superior capacity of PLNP to drive effector differentiation (**Fig. 4b**). These Runx2^+^ effector T cells displayed upregulated expression of genes involved in cytotoxicity (Gzmb, Prf1), effector functions (Cx3cr1, Tbx21), and differentiation and persistence (Zeb2, Runx2) (**Fig. S9a**), indicating robust induction of CD8^+^ T cell effector responses by PLNP. Additionally, Il7r^+^ effector T cells, associated with maintenance of memory T cells following vaccination, were more enriched in the PLNP group (∼25%) than in the LNP group (∼10%) (**Fig. 4b**), suggesting that PLNP promotes superior memory formation. To further elucidate the functional differences between LNP and PLNP vaccination, we analyzed differentially expressed genes (DEGs) between these two groups. Of the 2,303 DEGs identified (log fold change > 0.25), genes related to TCR signaling, effector functions, migration, and GTPase activation were significantly upregulated in the PLNP group (**Fig. 4c**). Gene set enrichment analysis (GSEA) further demonstrated elevated signatures for TCR signaling, effector functions, and migration in CD8^+^ T cells following PLNP vaccination (**Figs. 4d–f**). Furthermore, relative to LNP, PLNP resulted in greater enrichment of the IL12-STAT4 signaling pathway, indicating a more robust activation of CD8^+^ T cells through enhanced IL-12 production and downstream STAT4 signaling (**Fig. 4g**). The single-cell RNA-seq data of CD8^+^ T cells demonstrate that PLNP more effectively stimulates CD8^+^ T cells and enhances their effector functions compared to LNP.

**Figure 4.**
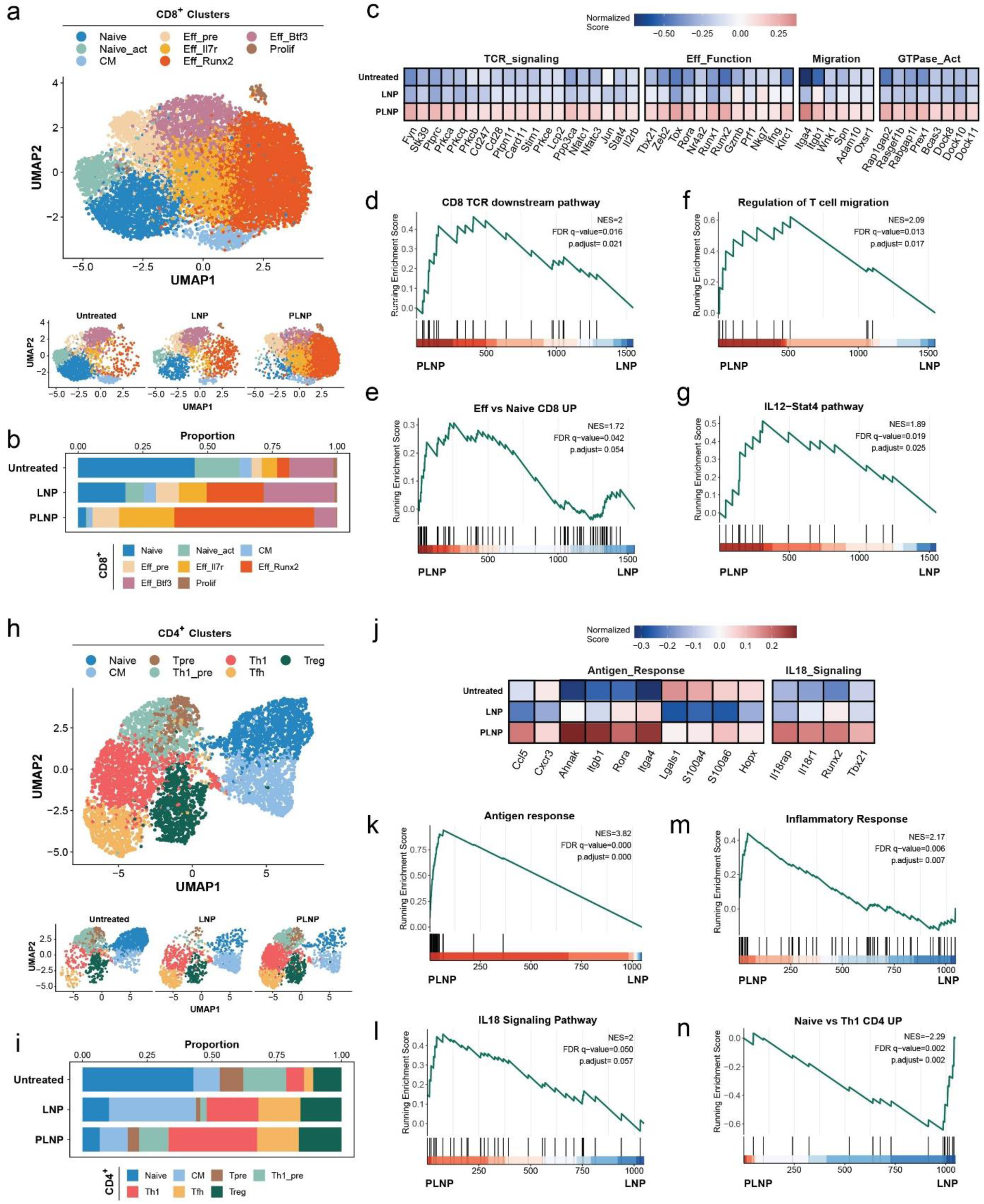
T-cell characterization by single-cell RNA sequencing following vaccination with LNP and PLNP. Mice were vaccinated with spike mRNA-loaded LNP and PLNP on day 0 and day 21 and splenocytes were collected on day 28 for characterization. Dose: 10 µg mRNA/mouse. n = 5. (a) Uniform Manifold Approximation and Projection (UMAP) depicting 8 distinct CD8^+^ T cell clusters, including naïve cells (Naïve), naïve cells with an early activation signature (Naïve_Act), central memory cells (CM), precursor effector cells (Eff_pre), effector cells expressing Il7r (Eff_Il7r), Runx2 (Eff_Runx2), and Btf3 (Eff_Btf3), and proliferative cells (Prolif). Top: overall UMAP of combining three conditions. Bottom: separate UMAP for each condition. (b) Stacked bar graph depicting proportions of each CD8^+^ T-cell subset comparing different groups. (c) Heatmap of top DEGs grouped by clusters related to cytotoxicity, effector, migration, and activation (GTPase). Color scale indicates mean expression in each group. (d-g) GSEA showing selected pathways enriched in CD8^+^ T cells in PLNP treated group versus LNP treated group. (h) UMAP depicting 7 distinct CD4^+^ T cell clusters, including naïve (Naïve), central memory (CM), precursors of both Th1 and Tfh (T_pre), precursors of Th1 (Th1_pre), T helper type 1 (Th1), T follicular helper (Tfh), and regulatory T cell (Treg). Top: overall UMAP of combining three conditions. Bottom: separate UMAP for each condition. (i) Stacked bar graph depicting proportions of each CD4^+^ T-cell subset comparing different groups. (j) Heatmap of top DEGs by clusters related to antigen response and IL-18 signaling. Color scale indicates mean expression in each group. (k-n) GSEA showing selected pathways enriched in CD4^+^ T cells in PLNP treated group versus LNP treated group. A positive enrichment score indicates enrichment in PLNP treated group versus LNP treated group, and a negative enrichment score indicates enrichment in LNP treated group versus PLNP treated group. NES: normalized enrichment score. The *P* value was generated using the permutation test and adjusted for multiple testing using the Benjamini-Hochberg procedure, and the FDR was estimated using the FDR q-values.

We next extended our transcriptomic analysis to CD4^+^ T cells. We identified seven distinct clusters representing key functional subsets, including naïve (Naïve), central memory (CM), precursors of both Th1 and Tfh (T_pre), precursors of Th1 (Th1_pre), T helper type 1 (Th1), T follicular helper (Tfh), and regulatory T cell (Treg) (**Figs. 4h and S9b**). Both mRNA-LNP and-PLNP vaccination shifted the distribution of these CD4^+^ T cell subsets when compared to untreated controls, with PLNP driving a more marked alteration in the overall repertoire. Notably, PLNP vaccination resulted in a significant enrichment of Th1 cells (∼30%), far exceeding the proportions seen in the LNP group (∼18%) (**Fig. 4i**). These Th1 cells upregulated genes associated with lineage commitment and effector functions, including Il18r1, Id2, Ccl5, Tbx21, and Nkg7 (**Fig. S9b**), indicative of potent Th1 polarization. In contrast, Tfh cell frequencies were comparable between the PLNP (∼14%) and LNP (∼15%) groups, suggesting equivalent capacity to drive B cell maturation and antibody production. We next examined DEGs between three different groups. Notably, genes associated with antigen response and genes in the IL-18 signaling pathway were all significantly upregulated in the PLNP group compared to both the LNP and untreated groups (**Fig. 4j**). These transcriptional changes mirror our *in vitro* findings, where PLNP strongly promoted Th1-biased cytokine secretion, with a notable increase in IL-12 production (**Fig. 1i**). As IL-12 plays an essential role in upregulating IL-18 receptor expression on Th1 cells, this leads to enhanced IL-18 signaling, promoting Th1 polarization and increased IFN-γ production. Consistent with these effects, GSEA confirmed that antigen response, IL-18 signaling, inflammatory response, and Th1 gene signatures were all more highly enriched in the PLNP group than in the LNP group (**Fig. 4k–n**). These results demonstrate that PLNP more effectively elicits Th1 polarization and supports robust cellular immunity.

Collectively, our single-cell RNA sequencing data show that mRNA-PLNP vaccination more effectively remodels both CD8^+^ and CD4^+^ T cell populations compared to traditional LNP formulations, leading to enhanced effector CD8^+^ T cell responses and stronger Th1 polarization. This dual action results in a more robust cellular immunity, which is critical for long-term protection and effective defense against infection and cancer.^23^

### mRNA-PLNP vaccine elicits strong humoral immunity

Having established the superiority of PLNP in cellular immunity, we next evaluated its humoral immune response. We assessed B cell activation, antibody production, antigen neutralization, and host protection, key parameters of humoral immunity.

To analyze B cell responses, C57BL/6 mice were immunized twice (day 0 and day 21) with spike mRNA-loaded LNP or PLNP (WT spike mRNA and B.1.1.529 spike mRNA, 1:1). Spleens and draining lymph nodes were collected on day 28 for flow cytometry analysis. PLNP vaccination resulted in higher frequencies of memory B cells and germinal center B cells in lymph nodes compared to LNP or untreated controls (**Figs. 5a-b, S10**), consistent with enhanced lymph node targeting by PLNP (**Fig. 2b**). Furthermore, PLNP induced greater frequencies of spike RBD-specific B cells and memory B cells in the spleen (**Figs. 5c-d, S11**).

**Figure 5.**
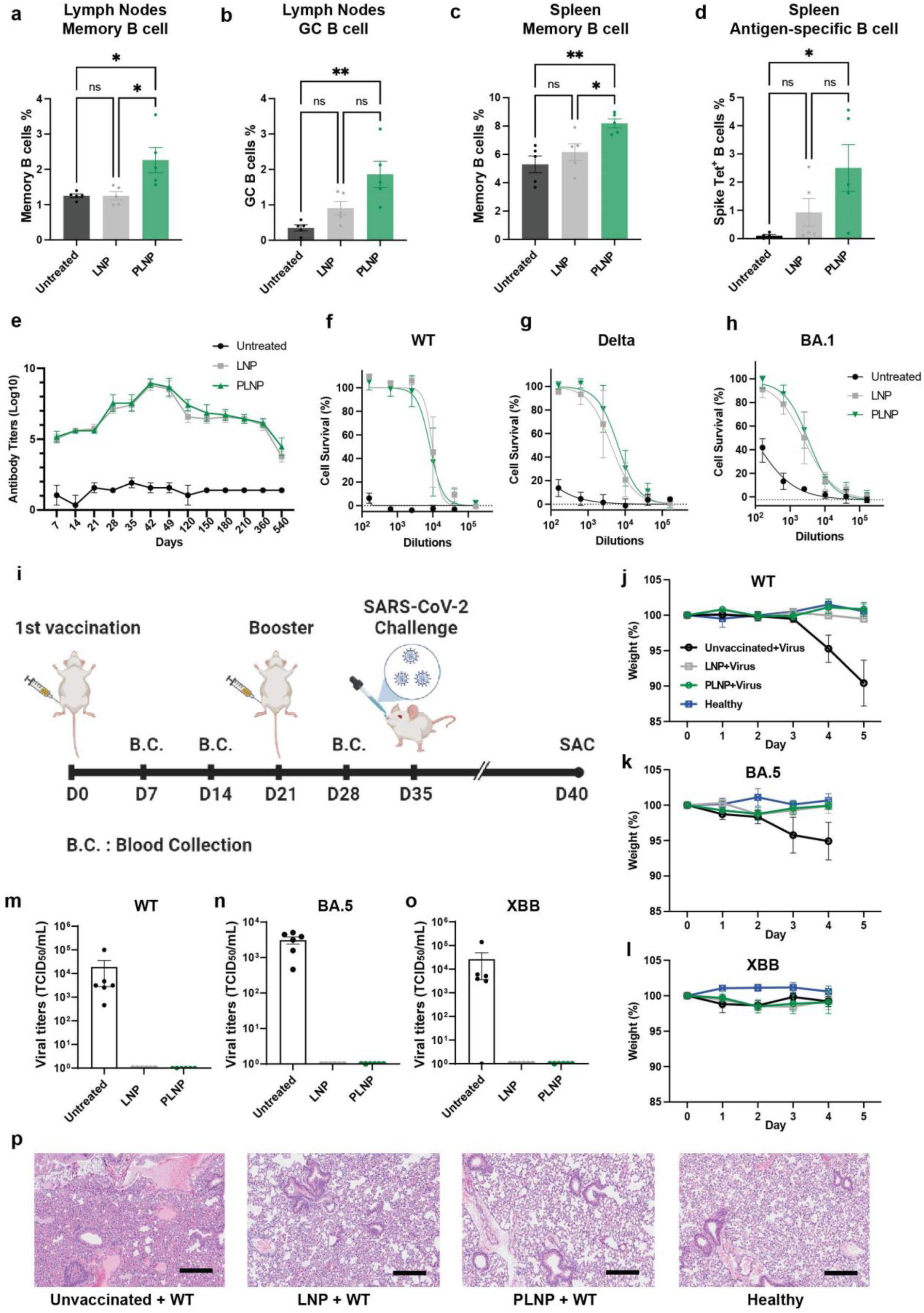
B cell response, antibody production, and protection against viral challenge post vaccination with LNP and PLNP. (a, b) B cell populations in lymph nodes collected on day 28. Mice were vaccinated with spike mRNA-loaded LNP and PLNP on day 0 and day 21. Dose: 10 µg mRNA/mouse. n = 5. (c, d) B cell populations in spleens collected on day 28. Mice were vaccinated with spike mRNA-loaded LNP and PLNP on day 0 and day 21. Dose: 10 µg mRNA/mouse. n = 5. (e) Spike RBD-specific antibody levels in serum samples collected from mice at different time points (day 7 to day 540). Mice were vaccinated with spike mRNA-loaded LNP and PLNP on day 0 and day 21. Dose: 10 µg mRNA/mouse. n = 5. (f-h) Neutralization of different SARS-CoV-2 variants (WT, Delta, and BA.1) by serum samples collected on day 35. Mice were vaccinated with spike mRNA-loaded LNP and PLNP on day 0 and day 21. Dose: 10 µg mRNA/mouse. n = 5. (i) Schematic illustration of SARS-CoV-2 challenge experiments. Mice were vaccinated with spike mRNA-loaded LNP or PLNP on day 0 and day 21 and further challenged with different SARS-CoV-2 variants (WT, BA.5 and XBB) on day 35. Dose: 10 µg mRNA/mouse. n = 6. (j-l) Mouse weight change after being challenged with different SARS-CoV-2 variants (WT, BA.5 and XBB). (m-o) TCID_50_ virial titers in mouse lungs after being challenged with different SARS-CoV-2 variants (WT, BA.5 and XBB). (p) Hematoxylin and Eosin (H&E)-stained lung tissues collected from mice challenged with SARS-CoV-2 WT virus. Scale bars: 50 µm. Data are presented as mean ± s.e.m. Statistical significance was determined using one-way ANOVA with Tukey post hoc test for multiple comparisons (a, b, c, d). ns: no significance, * P < 0.05, and ** P < 0.01.

Long-term antibody production was measured in C57BL/6 mice immunized with spike mRNA-loaded LNP or PLNP on day 0 and day 21, with serial blood collection up to 540 days post-immunization. Despite the differences in B cell numbers, PLNP and LNP generated comparable antibody titers throughout the 1.5-year period, with PLNP titers occasionally exceeding those of LNP (**Fig. 5e**). Similar results were seen in BALB/c mice vaccinated with influenza HA mRNA-loaded LNP or PLNP (**Fig. S12**).

We further performed neutralization assays using authentic (live) SARS-CoV-2 variants. Serum samples from vaccinated mice were incubated with diverse SARS-CoV-2 variants, followed by infection of Vero E6 cells. Neutralization efficiency was quantified by measuring the cell viability of Vero E6 cells after infection. Antibodies elicited by both LNP and PLNP formulations demonstrated similar neutralization capacity against WT, Delta, and BA.1 variants (**Figs. 5f-h**), indicating broad protection generated by both formulations.

### Prophylactic mRNA-PLNP vaccination protects against SARS-CoV-2 variants

We next examined whether the robust T and B cell responses induced by mRNA-PLNP vaccine can generate immune protection following SARS-CoV-2 infection. K18-hACE2 transgenic mice, which express human ACE2, were immunized twice on day 0 and day 21 with spike mRNA-loaded LNP or PLNP (WT spike mRNA and B.1.1.529 spike mRNA, 1:1) and subsequently challenged on day 35 with various SARS-CoV-2 variants, including WT, BA.5, and XBB (**Fig. 5i**). Healthy, untreated K18-hACE2 mice served as unvaccinated and uninfected controls.

Protection was assessed by monitoring body weight and animal activity post-infection. Unvaccinated mice infected with WT or BA.5 exhibited significant weight loss and elevated clinical scores (**Figs. 5j-k, S13a-b**). However, unvaccinated mice infected with XBB showed minimal weight loss (**Figs. 5l, S13c**), aligning with previous studies that report XBB’s decreased severity related to similar mutational effects.^49,50^ Importantly, all vaccinated mice, regardless of LNP or PLNP formulation, were protected from weight loss (**Figs. 5j-l**) across all challenged variants. This demonstrates strong and broad protective efficacy by both PLNP and LNP.

Because XBB infections did not significantly affect body weight, further evaluation was carried out. Five (WT) or four (BA.5 or XBB) days post-challenge, mice were sacrificed, and lungs, a primary site of infection, were collected and homogenized for quantification of acute viral loads. High levels of viral titers were detected in unvaccinated mice infected with WT, BA.5, and XBB, while no detectable virus was present in the lungs of vaccinated mice (**Figs. 5m-o**). In addition, all vaccinated mice were protected from lung tissue damage after WT viral challenge, while unvaccinated mice showed pronounced tissue damage (**Fig. 5p**). These findings confirm that the mRNA-PLNP vaccine confers effective and broad protection against SARS-CoV-2 variant infection.

### Therapeutic mRNA-PLNP exhibits potent anti-tumor efficacy in solid tumor models

At first glance, the studies above may suggest that the protective efficacy of LNP and PLNP is comparable, but it is important to clarify that the SARS-CoV-2 challenge mouse model is not stringent enough to discern differences in vaccine efficacy. Even a single shot of a spike vaccine elicits robust immune responses that mediate near complete protection in this model, rendering that it is inadequate for discerning differences in vaccine efficacies^51–53^ Therefore, to further elucidate and differentiate the functional advantages of PLNP over LNP, subsequent investigations were performed in a solid tumor model, where more subtle differences in immunological efficacy could be resolved.

We first utilized B16-OVA melanoma mouse model to evaluate the anti-tumor efficacy of PLNP. OVA mRNA-loaded LNP or PLNP were prepared, and therapeutic interventions were conducted at distinct tumor stages, early-stage (∼30 mm³) and advanced-stage (∼85 mm³), to assess the relative efficacy of PLNP compared to LNP in the treatment of solid tumors.

For early-stage tumors, C57BL/6 mice were inoculated with 0.5 × 10⁶ B16-OVA cells on day 0. Upon tumors reaching ∼30 mm³ on day 7, the mice received OVA mRNA-loaded LNP or PLNP treatment, followed by a booster dose on day 10 via intramuscular injection (**Fig. 6a**). Both therapeutic vaccines significantly reduced tumor burden, completely eradicating tumors by day 16with 100% survival, whereas all untreated mice exhibited rapid tumor progression and succumbed prior to day 23. To assess the induction of immunological memory, LNP or PLNP-treated mice were rechallenged with B16-OVA cells on day 25. Notably, both LNP- and PLNP-treated groups remained tumor-free (**Figs. 6b-c**), indicating both mRNA vaccine formulations established durable T cell memory capable of preventing tumor recurrence.

**Figure 6.**
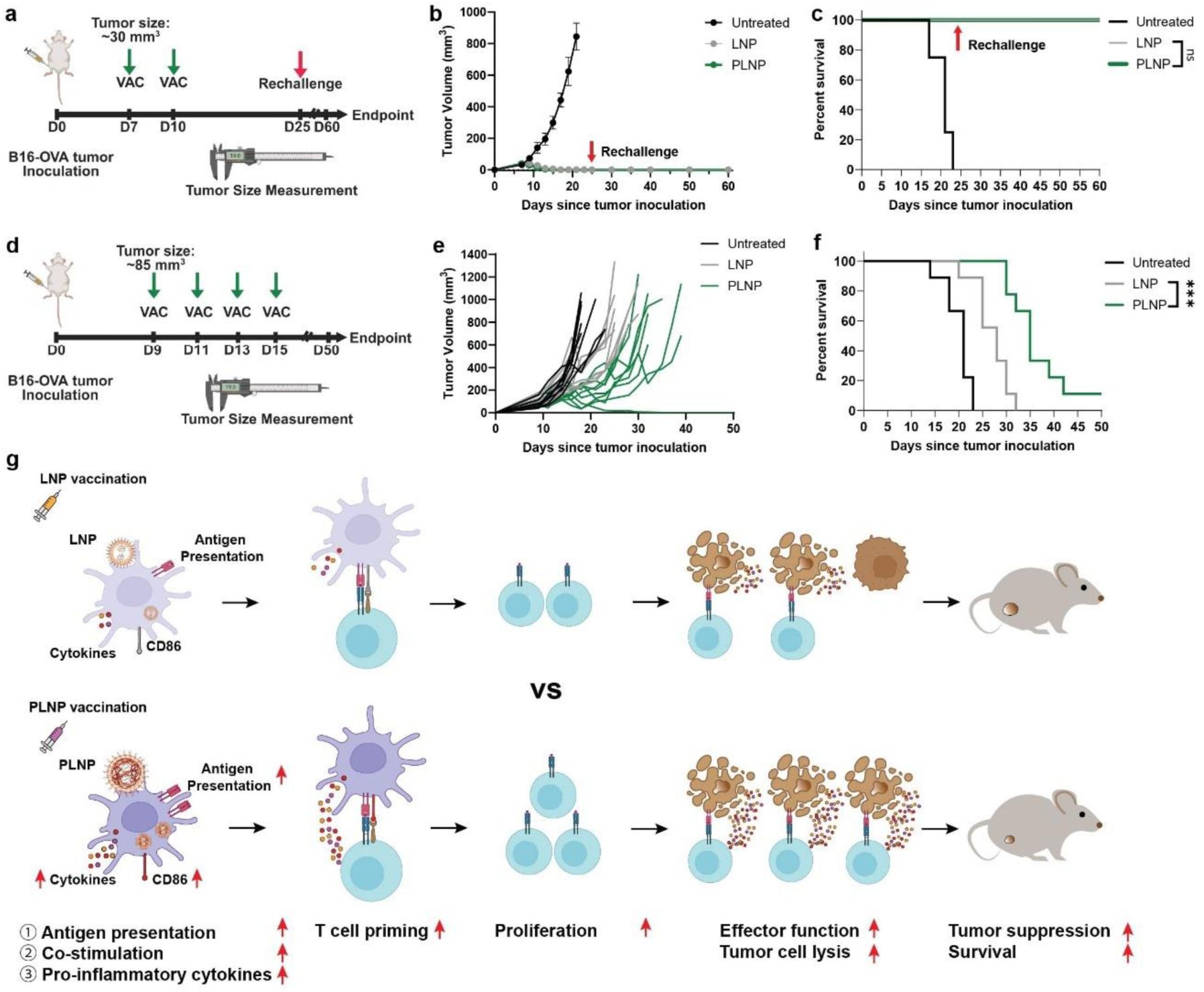
PLNP generates potent anti-tumor efficacy for treating melanoma. (a) Schematic illustration of B16-OVA melanoma tumor inoculation (day 0), treatment (day 7 and day 10), and rechallenge (day 25) timeline (early-stage tumor model). (b) Tumor size post treatment with OVA mRNA-loaded LNP or PLNP on day 7 and day 10. Dose: 5 µg mRNA/mouse. n = 4. Data are presented as mean ± s.e.m. (c) Mouse survival after treatment with OVA mRNA-loaded LNP or PLNP on day 7 and day 10 and after rechallenging on day 25. (d) Schematic illustration of B16-OVA melanoma tumor inoculation (day 0) and treatment (day 9, day 11, day 13, and day 15) (advanced-stage tumor model). (e) Tumor size post treatment with OVA mRNA-loaded LNP or PLNP on day 9, day 11, day 13, and day 15. Dose: 5 µg mRNA/mouse. n = 9. (f) Mouse survival after treatment with OVA mRNA loaded LNP or PLNP on day 9, day 11, day 13, and day 15. Statistical significance was determined using Holm-Šídák’s multiple comparisons test. *** P < 0.001. (g) Schematic illustration of PLNP enhancing antigen-specific T cell response for generating better anti-tumor efficacy.

Finally, we increased the stringency of the tumor model using an advanced-stage tumor model, in which treatment was initiated when tumors reached ∼85 mm³ on day 9. Mice received OVA mRNA-loaded LNP or PLNP, followed by three booster injections on days 11, 13, and 15 (**Fig. 6d**). In this setting, mRNA-PLNP vaccination showed markedly superior tumor control and extended survival compared to the LNP-treated cohort (**Figs. 6e-f**), demonstrating its enhanced therapeutic efficacy against established solid tumors.

These results demonstrate that therapeutic mRNA-PLNP vaccines effectively induce potent T cell-mediated anti-tumor immunity, with particularly notable benefits in the management of larger solid tumors. The enhanced therapeutic efficacy of PLNP compared to LNP is likely associated with superior antigen presentation, co-stimulation, and cytokine expression, which are three critical signals that are mechanistically required for the activation of T cell responses (**Fig. 6g**).

## DISCUSSION

The advent of COVID-19 mRNA vaccines has fundamentally transformed approaches to the development of prophylactic vaccines for infectious diseases.^1,2^ Historically, the primary endpoint for evaluating vaccine efficacy has centered on the induction of neutralizing antibodies, given their critical role in conferring protective immunity. In response to this emphasis, new LNP formulations have been developed to enhance the production of neutralizing antibodies, further advancing the effectiveness of mRNA vaccine platforms in inducing robust responses.^54,55^ However, accumulating evidence demonstrates that SARS-CoV-2 variants possess the capacity to evade antibody-mediated neutralization.^56,57^ Consequently, T cell-mediated immunity has gained attention as a crucial component for controlling newly emerging variants, since T cells can recognize conserved linear epitopes.^58,59^ Parallel advances have extended the applications of therapeutic mRNA vaccines to oncology, where robust anti-tumor responses mediated by T cells are imperative for tumor eradication.^27,60^ Recent studies have emphasized that the generation of targeted T cell responses is essential for effective cancer immunotherapy,^24,61^ as well as for pandemic preparedness against rapidly evolving pathogens.^17,19^ These findings underscore the urgent need for next-generation mRNA vaccines that can elicit robust T cell-mediated immunity in addition to strong humoral responses.

In this study, we engineered a novel PLNP mRNA delivery platform designed specifically to enhance T cell-mediated cellular immunity while preserving potent B cell-mediated humoral immunity. This novel PLNP formulation markedly improved mRNA translation and lymph-node targeting, which in turn enhanced antigen presentation, APC co-stimulation, and pro-inflammatory cytokine secretion, three essential signals for effective T-cell priming. The coordinated augmentation of these signals drove robust antigen-specific T-cell responses, ultimately resulting in improved tumor control. Importantly, this platform substantially augmented cellular immunity without compromising the humoral response and maintained effective antibody production, which is likely attributable to its efficient lymph node targeting and the resultant broad immune activation of B cells.^62,63^ Achieving this balanced immune profile is essential for next-generation vaccine platforms, which must be capable of inducing both robust cellular and humoral immune responses to deliver comprehensive and durable protection tailored to the specific immunological demands of diverse diseases.^28,36^

Our PLNP formulation, in principle, is compatible with a broad range of emerging mRNA technologies, including circular mRNA,^64^ self-amplifying mRNA,^65^ branched mRNA,^66^ capped mRNA,^67^ modular mRNA,^68^ and sequence-optimized mRNAs through viral sequence insertion^69^ or deep generative modeling.^70,71^ When used in conjunction with innovative mRNA design strategies, PLNP-mediated delivery can synergistically enhance immune activation by promoting efficient lymphoid targeting, APC uptake and activation, and robust antigen presentation. Such a combined approach holds the potential to maximize both the magnitude and durability of T-cell immunity, offering a scientific rationale for the development of next-generation mRNA vaccines with superior breadth, efficacy, and adaptability for emerging pathogens and complex diseases.

Current personalized cancer vaccine strategies focus on identifying and engineering mRNA sequences that encode patient-specific tumor neoantigens, thereby optimizing antigen structure to improve T cell recognition and activation.^72,73^ Advances in mRNA sequence design, including AI-driven epitope prediction^74,75^ and codon optimization,^76,71^ have shown promise for optimizing cancer vaccines. However, achieving clinical efficacy increasingly requires the integration of these molecular innovations with stable and immunogenic delivery platforms. The PLNP system provides a crucial adjunct, as its advanced delivery capabilities further amplify T cell activation and antitumor immunity when used in combination with personalized neoantigen mRNA constructs.^72,77^ This synergistic approach holds the potential to address key challenges such as tumor heterogeneity^78^ and limited immune engagement^79^. Therefore, the convergence of rational mRNA sequence engineering with optimized PLNP delivery represents a promising paradigm for eliciting robust and durable antitumor T cell responses in individualized cancer vaccine regimens.

The unique properties of PLNPs also position them as promising delivery vehicles for *in vivo* cell therapy applications.^80–82^ Conventional LNP-based systems have proven highly effective for delivering mRNA into target cells and tissues, overcoming major barriers such as instability and degradation of nucleic acids *in vivo*.^3,37^ By encapsulating and protecting mRNA cargo, PLNPs can facilitate efficient translation and expression of therapeutic genes directly within recipient cells, thereby enabling advanced cell therapy strategies. This includes the transient reprogramming of immune cells or somatic cells *in vivo*, for applications such as CAR-T cell engineering,^80^ immune system modulation,^3^ and tissue regeneration.^82^ In addition, PLNPs can be tailored for cell-specific targeting, potentially through modification with targeting ligands or peptides that ensure selective delivery to desired cell populations and thereby minimize off-target effects.^83^ Overall, the adaptability and efficiency of PLNPs as mRNA delivery platforms hold significant promise for expanding the scope and impact of *in vivo* cell therapies for cancer and other challenging diseases.

Despite significant advances in our PLNP formulation, further improvements in biodistribution and targeting specificity remain necessary for optimized therapeutic outcomes.^83,84^ Although PLNPs markedly enhance *in vivo* mRNA translation, our studies revealed pronounced expression in the liver, attributable to uptake by resident Kupffer cells. This off-target hepatic accumulation potentially diminishes the efficiency of mRNA delivery to desired tissues and may introduce unwanted side effects. Addressing organ-specific targeting requires rational adjustments to nanoparticle composition, such as the incorporation of alternative helper lipids, ionizable lipids, and specific surface components, or modifying physicochemical properties, all of which can decrease non-specific liver uptake, promote selective tissue accumulation^85^, reduce systemic toxicity^86^, and fine-tune immune activation^54^. Advanced biodegradable polymers like poly(β-amino ester) and poly(amine-co-ester) are also under investigation for their promising biocompatibility and controlled release profiles.^87,88^ In parallel, cell-specific delivery within targeted organs is essential to maximize efficacy and minimize off-target actions.^89^ This can be achieved through active targeting strategies, such as functionalizing PLNP surfaces with ligands or antibodies directed against cell-specific markers, thereby driving selective uptake by APCs in lymphoid tissues or tumor-infiltrating immune cells.^83,84^ Continued optimization focused on both organ-level and cell-level targeting will be essential to fully realize the clinical potential of PLNP-mediated mRNA delivery platforms.

In summary, our findings demonstrate that the rationally engineered PLNP-based mRNA vaccine platform enables the effective integration of potent T cell priming with robust humoral immunity. By optimizing mRNA delivery into APCs, enhancing mRNA translation efficiency, facilitating precise lymph node targeting, and promoting strong antigen-specific CD8^+^ T cell responses along with Th1 polarization, this advanced formulation addresses critical bottlenecks encountered by conventional mRNA vaccines. The balanced induction of both cellular and antibody-mediated immune responses underscores the versatility and clinical potential of our platform. Importantly, the modular nature of PLNPs permits broad compatibility with emerging mRNA technologies and may support the development of highly personalized immunotherapies. Altogether, our strategy offers a promising and adaptable approach for manufacturing next-generation mRNA therapeutics designed to target a diverse spectrum of indications, ranging from infectious diseases to solid tumors and other challenging clinical contexts where robust T cell-driven immunity is essential for lasting protection and therapeutic success.

## METHODS

### mRNA synthesis

Wild-type SARS-CoV-2 spike encoding plasmid was obtained from the MacMaster lab. Omicron spike (B.1.1.529), influenza A H1N1 (A/Puerto Rico/8/1934) hemagglutinin (HA), and FLuc sequences were obtained from commercially available omicron spike, HA, and FLuc Gene ORF cDNA clone expression plasmids (Sino Biological #VG40835-UT, Sino Biological #VG11684-UT, and Addgene #101156) and inserted into the same plasmid backbone using Gibson assembly. Plasmids containing the T7 promoter were transformed into E. coli for amplification, and plasmid DNAs were subsequently purified using the Qiagen Midiprep Kit (Qiagen #12123). Antigen mRNAs were synthesized by *in vitro* transcription using the HiScribe® T7 High Yield RNA Synthesis Kit (NEB #E2040L) with N1-methylpseudouridine (TriLink #N-1081) modification. The transcribed mRNAs were subsequently capped with the Vaccinia Capping System (NEB #M2080S) and mRNA Cap 2’-O-Methyltransferase (NEB #M0366), followed by poly (A) tailing using E. coli Poly(A) Polymerase (NEB #M0276). The resulting mRNAs were purified using the Monarch^®^ RNA Cleanup Kit (NEB #T2050), and the integrity and purity of the mRNA were verified by gel electrophoresis. OVA mRNA was purchased from GenScript (#SC2346).

### Lipid nanoparticle preparation

The LNP formulation consisted of ionizable lipid, helper lipid, cholesterol, and PEG-lipid at a weight ratio of 57.2:12.7:24.0:6.0. Lipids were dissolved in ethanol, while mRNAs were dissolved in 10 mM citrate buffer (pH 4.5). PLNP were formulated similarly, with the addition of the biocompatible polymer poly(lactic-co-glycolic acid (PLGA). The weight ratio of ionizable lipid, helper lipid, cholesterol, PEG-lipid, and PLGA is 47.1:10.5:19.8:5.0:17.7. DSPC was dissolved in ethanol and further mixed with other lipids and polymer dissolved in acetone. mRNAs were dissolved in 10 mM citrate buffer (pH 4.5). LNP and PLNP were prepared by rapidly mixing the aqueous phase (mRNAs) with the organic phase (lipids and polymer) using a microfluidic mixer (NanoAssemblr™ Ignite System). Following formulation, excess solvent was removed by centrifugal filtration (MWCO:100 kDa, 4000 rpm). The resulting nanoparticles were washed three times with 1×PBS and sterilized by passing through a 0.2 µm filter.

### Characterization of the lipid nanoparticles

The hydrodynamic diameter and polydispersity index of LNP and PLNP were measured by dynamic light scattering (DLS) using a Zetasizer Nano-S (Malvern). The morphologies of LNP and PLNP were imaged by transmission electron microscopy (JEM-2010F, JEOL, Japan). The mRNA encapsulation efficiency was determined using the QuantiFluor® RNA System (Promega #E3310) following lysis of the nanoparticles with 0.1% Triton-X100.

### *In vitro* characterization of mRNA translation and cytokine generation in macrophages

RAW 264.7 cells (3 × 10^5^ cells per well) were seeded in 6-well plates and cultured in cell culture medium (DMEM supplemented with 10% FBS and 1% penicillin/streptomycin) at 37°C in a humid incubator with 5% CO_2_ for 24 h. LNP or PLNP encapsulating spike mRNAs (wild-type and omicron B.1.1.529, 1:1, 1 µg/mL) was added to 6-well plate and incubated for 24 hours (cytokine characterization) or 48 hours (mRNA translation characterization). For cytokine characterization, intracellular cytokine staining was performed using anti-TNF-α (BV605), anti-IFN-γ (PE/Cy7), and anti-IL-2 (BV650). Samples were analyzed on a BD LSRFortessa flow cytometer and data were processed with FlowJo software. For mRNA translation characterization, cells were washed with 1 × PBS and stained with LIVE/Dead™ Fixable Violet viability dye and anti-spike AF647 antibody (eBioscience). The mRNA translation was analyzed using a BD LSRFortessa flow cytometer and data were processed with FlowJo software. To monitor mRNA translation at different time points, RAW 264.7 cells (3 × 10^4^ cells per well) were seeded in 6-well plates and cultured in cell culture medium (DMEM supplemented with 10% FBS and 1% penicillin/streptomycin) at 37°C in a humid incubator with 5% CO_2_. LNP or PLNP encapsulating FLuc mRNA (1 µg/mL) was added to the plates. Cells were harvested on day 1, 2, 3, or 5 and plated into 96-well plates to quantify the FLuc expression by ONE-Glo™ FLuc Assay System (Promega #E6110) and microplate reader (BioTek) and the signals were normalized to day-1 signal of LNP-treated group.

### *In vitro* characterization of antigen presentation

Bone marrow-derived dendritic cells were collected from C57BL/6 female mice and cultured in RPMI 1640 medium (10% FBS, 1% penicillin/streptomycin, 2mM L-glutamine, and 0.05mM 2-mercaptoethanol) supplemented with granulocyte-macrophage colony-stimulating factor (GM-CSF, 20 ng/mL).^90^ The cells were plated in 6-well plates (1 × 10^6^ cells per well) and incubated with OVA mRNA-loaded LNP or PLNP (1 µg/mL) for 48 hours. Cells were then washed with 1 × PBS and stained with LIVE/Dead™ Fixable Violet, anti-CD45 (bBV605), anti-CD11c (PE-Cy7), and anti-mouse H-2Kb bound to SIINFEKL (PE). The translationantigen presentation was analyzed using a spectral flow cytometer (Cytek Aurora) and data were processed with FlowJo software.

### *In vitro* human/mouse PBMC stimulation inflammatory cytokine measurement

Human or mouse peripheral blood mononuclear cells (PBMCs; 1×10^6^ cells per well) were thawed and cultured in RPMI 1640 medium supplemented with 10% FBS, 1% penicillin/streptomycin, 2mM L-glutamine, and 0.05mM 2-mercaptoethanol. Different formulations of empty LNP, PLNP, spike mRNA-loaded LNP or spike mRNA-loaded PLNP (wild-type and omicron B.1.1.529; 1:1; 1 µg/mL) were added to each well. After incubation for 24 h at 37 °C in a humidified 5% CO₂ incubator, cell culture supernatants were collected by centrifugation at 300 g for 5 min. Inflammatory cytokine panels were quantified using the LEGENDplex Human or Mouse Inflammation Panel (Biolegend) according to the manufacturer’s instructions. Samples were acquired on a spectral flow cytometer (Cytek Aurora) and analyzed using the LEGENDplex Data Analysis Software Suite.

### Animals

C57BL/6 female mice, BALB/c female mice, and B6.Cg-Tg(K18-ACE2)2Prlmn/J (K18-hACE2) male and female mice at the age of 6-8 weeks old were purchased from Jackson Lab and maintained in the Animal Facility of the University of Chicago. All animal experiments were approved by the Institutional Animal Care and Use Committee (IACUC) of the University of Chicago.

### *In vivo* mRNA translation and lymph node targeting

For bioluminescent imaging to track the FLuc mRNA translation *in vivo*, BALB/c mice were intramuscularly injected with LNP or PLNP encapsulating FLuc mRNA (5 µg/mouse, n = 5). At 4, 24, 48, 72, and 96 h post injection, luciferin substrate (freshly reconstituted with 1×PBS) was administered intraperitoneally. Mice were anesthetized and imaged using Spectral Lago X under default acquisition settings. The bioluminescent signals were quantified with Aura 4.0 imaging software. Mice injected with 1×PBS as negative control and background luminescence from these controls were subtracted from all measurements.

For fluorescent imaging of the draining lymph nodes, C57BL/6 mice were intramuscularly injected with DiD-labelled spike mRNA-loaded LNP or PLNP (10 µg mRNA/mouse, n = 4). At 24 h post injection, draining lymph nodes were collected and imaged by Spectral Lago X under default acquisition settings. Mice injected with 1×PBS served as negative controls and background fluorescent from these controls was subtracted from experimental measurements.

### Characterization of innate immune cell activation and systematic inflammatory response

Draining lymph nodes were collected from C57BL/6 mice on day 28 post vaccination with spike mRNA-loaded LNP or PLNP (day 0 and day 21, 10 µg mRNA/mouse, n = 5) and transferred into 24 well plates containing digestion medium composed of DMEM supplemented with 1% penicillin–streptomycin, 2% FBS, 1.2 mM CaCl_2_, collagenase D (1:50) and collagenase IV (1:100). Plates were sealed with parafilm and aluminum foil, sprayed with ethanol for sterilization, and incubated at 37 °C for 45 min in a shaking incubator. Following digestion, lymph nodes were transferred onto 70 µm cell strainers placed over 50 mL conical tubes and gently dissociated using a 1 mL syringe plunger. The strainers were rinsed with DMEM containing 2% FBS to a total volume of 20 mL. Cell suspensions were centrifuged at 1,500 rpm for 8 min, washed once with 5 mL complete DMEM medium, and resuspended in 0.5 mL complete DMEM medium. Single cell suspensions were stained with Live/dead Zombie UV™, anti-PDCA-1 (BUV563), anti-F4/80 (BUV737), anti-CD8a (BUV805), anti-CD11c (BV421), anti-NK1.1 (BV510), anti-CD45 (BV605), anti-CD11b (BV650), anti-CD80 (BV711), anti-Ly6C (BV785), anti-CD3 (AF488), anti-CD19 (PerCP-Cy5.5), anti-CD103 (PE), anti-Siglec-F (PE-CF594), anti-CD169 (PE-Cy7), anti-CD86 (AF647), anti-MHCII (AF700), and anti-Ly6G (APC-Cy7). Samples were analyzed on a spectral flow cytometer (Cytek Aurora) and data was processed using FlowJo software.

Blood samples were collected from C57BL/6 mice via submandibular vein puncture using a 4 mm sterile lancet at 24 hour post vaccination with spike mRNA-loaded LNP or PLNP (10 µg mRNA/mouse, n = 5). Approximately 100 µL of blood was collected into EDTA-coated heparinized tubes. The samples were centrifuged at 1,000 × g for 15 min at 4°C and the serum were collected by aspirating the supernatant. Inflammatory cytokine panels were quantified using the LEGENDplex Mouse Inflammation Panel (Biolegend) according to the manufacturer’s instructions. Samples were acquired on a spectral flow cytometer (Cytek Aurora) and analyzed using the LEGENDplex Data Analysis Software Suite.

### Characterization of T cell response

To access T cell responses following vaccination, groups of C57BL/6 mice (for spike mRNA-loaded LNP or PLNP, n = 5, 10 µg mRNA/mouse ; for OVA mRNA-loaded LNP or PLNP, n = 5, 5 µg mRNA/mouse) and BALB/c mice (for HA mRNA loaded-LNP or PLNP, 10 µg mRNA/mouse, n = 5) were intramuscularly injected with LNP or PLNP using a 29½G insulin syringe (BD Biosciences) on day 0 and day 21, respectively. Mice were euthanized on day 28 and spleens were harvested and mechanically dissociated into single cell suspension by pressing through 70 µm strainers using syringe plungers. Cells were washed with T cell medium (RPMI 1640 supplemented with 10% FBS, 1% penicillin/streptomycin, 2mM L-glutamine, and 0.05mM 2-mercaptoethanol) and divided into different fractions for different T cell analysis as noted below. To assess antigen-specific cytokine production by T cells, 1×10^6^ splenocytes were incubated with spike, HA, or OVA peptide pools (2 µg/mL) for 6 h at 37 °C in a humidified incubator with 5% CO₂. The three peptide pools are a pool 316 peptides derived from a peptide scan (15-mer sequences with 11 amino acids overlap) spanning the full-length spike glycoprotein (Genscript, RP30020), a pool of 139 peptides derived from a peptide scan (15-mer sequences with 11 amino acids overlap) through HA of influenza A H1N1 A/Puerto Rico/8/1934 (Pepmix #PM-INFA-HAPR), and a pool of lyophilized peptides derived from a peptide scan (15-mer sequences with 11 amino acids overlap) covering the complete sequence of chicken OVA (PepTivator # 130-099-771). Cell culture supernatants were collected after 6-hour stimulation, and cytokine secretion levels were measured using the LEGENDplex assay (BioLegend) according to the manufacturer’s protocol. For phenotypic analysis, 1 × 10⁶ splenocytes were stained with LIVE/Dead™ Fixable Violet, anti-TCR (PE), anti-CD4 (BUV496), anti-CD8 (FITC), and spike tetramer-AF647 (assembled at a 4:1 molar ratio using biotinylated major histocompatibility complex MHC monomer obtained from the NIH Tetramer Core Facility and streptavidin-AF647). For spike H2-Kb VNFNFNGL tetramer and OVA H2-Kb SIINFEKL tetramer, the cells were stained under 4 ⁰C for 30 min. For HA H2-Kd IYSTVASSL tetramer, the cells were stained under 37 ⁰C for 30 min. Samples were analyzed on a BD LSRFortessa flow cytometer to quantify antigen-specific T cell populations.

### Single-cell RNA sequencing

Following immunization on day 0 and day 21 with spike mRNA-loaded LNP or PLNP, mouse splenocytes were collected on day 28 for single-cell transcriptomic profiling of CD4^+^ and CD8^+^ T cell responses. Cryopreserved splenocytes were thawed and washed thrice with cold cell media (RPMI 1640, 10% FBS) prior to fluorescence-activated cell sorting (FACS). Cells were stained with Zombie NIR, anti-CD3 (BUV395), anti-CD4 (PerCP-Cy5.5) and anti-CD8 (FITC). CD4^+^ and CD8^+^ T cells were sorted separately and used for single cell RNA sequencing. Approximately10,000 live CD8^+^ T cells (CD3^+^CD8^+^) per sample encapsulated into droplets using the Chromium Next GEM Single-Cell 5’ Kit v2 (10x Genomics #1000263). cDNA libraries were prepared for Single-cell RNA sequencing. All libraries (RNA-seq) were quantified using the Qubit dsDNA HS Assay Kit (Invitrogen, Q32851) and assessed for fragment sizes with high-sensitivity D5000 ScreenTapes (Agilent, 5067-5592). Pooled libraries were sequenced on an Illumina Novaseq-6000 platform. Sequencing data were used to evaluate T cell expansion and functional phenotypes associated with LNP and PLNP vaccination.

### Characterization of B cell response

Draining lymph nodes and spleens were collected on day 28 post vaccination from C57BL/6 mice vaccinated with spike mRNA-loaded LNP or PLNP (day 0 and day 21, 10 µg mRNA/mouse, n = 5). Draining lymph nodes and spleens were processed to make single-cell suspensions. Red blood cells from the spleens were lysed by 1× RBC lysis buffer (eBioscience #00-4333-57). The spike RBD tetramers were prepared by mixing SARS-CoV-2 spike RBD His biotin protein (R&D #BT10500-050) with either streptavidin-PE or streptavidin-AF647 at a molar ratio of 4:1 and incubating for 30 min at 4°C. To characterize the B cell responses in the lymph nodes, single-cell samples were stained with LIVE/Dead™ Fixable Violet, anti-CD3 (FITC), anti-B220 (BUV737), anti-CD19 (BUV395), anti-CD38 (APC-Cy7), anti-GL7 (PerCP-Cy5.5), anti-IgD (BV605), anti-IgM (BV786), anti-CD138 (PE-Cy7), and spike RBD tetramer (PE and AF647). To characterize the B cell responses in the spleens, single-cell samples were stained with LIVE/Dead™ Fixable Violet, anti-CD3 (FITC), anti-CD19 (BUV395), anti-CD38 (PE-Cy7), anti-GL7 (PerCP-Cy5.5), anti-IgD (BV605), and spike RBD tetramer (PE and AF647) for 30 min at 4°C. The flow cytometry analysis was performed on a BD LSRFortessa and the data was analyzed by FlowJo.

### Antibody titer measurement by ELISA

Blood samples were collected after initial immunization for quantification of SARS-CoV-2 spike-specific neutralizing antibody (C57BL/6 mice vaccinated with spike mRNA-loaded LNP or PLNP, day 7 to day 540) or HA-specific neutralizing antibody (BALB/c mice vaccinated with HA mRNA loaded-LNP or PLNP, day 7, day 14, day 21, and day 28). Blood samples were collected via submandibular vein puncture using a 4 mm sterile lancet. Approximately 100 µL of blood was collected into EDTA-coated heparinized tubes. The samples were centrifuged at 1,000 × g for 15 min at 4°C and the serum were collected by aspirating the supernatant. 96-well ELISA plates (Nunc™ MaxiSorp™) were coated with SARS-CoV-2 spike proteins (obtained from the MacMaster lab) or influenza HA proteins (Sino Biological #11684-V08B) (1 µg/mL, 100 µL) and incubated overnight at 4°C. The plates were washed gently with 200 uL 1×PBS+0.05% Tween 20 for three times and blocked by adding 200 µL 1×Assay Diluent (BioLegend #421203) into each well. The plates were washed gently with 200 uL 1×PBS+0.05% Tween 20 for three times after 2 h incubation with 300 rpm shaking under room temperature. Serial diluted serum samples were added into the plates. After 1 h incubation with 300 rpm shaking under room temperature, the plates were washed gently with 200 uL 1×PBS+0.05% Tween 20 for three times. Anti-Mouse IgG-HRP antibody (SouthernBiotech #1030-05) (1000x dilution in 1x Assay Diluent, 100 µL) were added into the plates. After 1 h incubation with 300 rpm shaking under room temperature, the plates were washed gently 5 times using 200 µL 1×PBS+0.05% Tween 20. TMB substrate (100 µl, BioLegend #421101) were further added into each well and incubate for 20 min under dark. Finally, 100 μL stop reagent (2N H_2_SO_4_) was added into each well. The absorbance values at 450 nm were read by a microplate reader (Genios Tecan). Antibody endpoint titers were determined by establishing the threshold based on negative controls without adding any serum samples.

### Neutralization efficiency against SARS-CoV-2 variants

Vero E6 cells were seeded in 96-well plates at 2.5 × 10^5^ cells/mL (100 μl/well) and incubated at 37°C in a humid incubator with 5% CO_2_ for 24 h. Serum samples collected on D35 post vaccination were heat inactivated at 56°C for 30 minutes and serial diluted in sterile 96-well plates. A 75 µL of viral suspension containing 100 TCID_50_ of SARS-CoV-2 (WT, Delta, Omicron BA.1) was added to each serum dilution. After incubation for 1 hour at 37°C, 100 µL of the serum-virus mixture was transferred to Vero E6 cell plates and incubated for an additional 1 hour at 37°C. The inoculation was removed and replaced with fresh culture medium. After 24 h of incubation at 37°C, cells were stained by crystal violet staining and cell viabilities were measured by microplate reader (Tecan). Neutralization efficiency was determined based on the cell survival in groups infected by serum-treated virus relative to groups infected by virus-only.

### *In vivo* challenge with SARS-CoV-2 variants

All experiments involving live SARS-CoV-2 were conducted in biosafety level 3 (BSL3) and animal BSL3 (ABSL3) facilities at the University of Chicago, following institutional guidelines and protocols approved by the Institutional Biosafety Committee (IBC) and the Institutional Animal Care and Use Committee (IACUC) at the University of Chicago. 6-8 weeks old male (n = 3) and female (n = 3) B6.Cg-Tg(K18-ACE2)2Prlmn/J (K18-hACE2) mice (Jackson Laboratory) were immunized with spike mRNA-loaded LNP or PLNP (10 µg/mouse) at day 0 and day 21. The mice were anesthetized by intraperitoneal injection with ketamine–xylazine (100 mg–20 mg/kg), for intranasal administration of virus on day 35. Animals were challenged with 2 × 10^4^ PFU of WT, 1 × 10^5^ of BA.5, or 1 × 10^5^ of XBB. Mice were monitored twice daily to record clinical symptoms and weighed daily post-challenge with virus. Categories in clinical scoring included: Score 0 (pre-inoculation) - animal is bright, alert, active, with normal fur coat and posture; Score 1 (post-inoculation, pi) – animal is bright, alert, active, normal fur coat and posture, no weight loss; Score 1.5 - animal has slightly ruffled fur but is active; weight loss under 2.5%; Score 2 (pi) – animal has ruffled fur, is less active; weight loss under 5%; Score 2.5 (pi) - animal has ruffled fur, is not active but moves when touched, may have hunched posture or difficulty breathing; weight loss 5-10%; Score 3 (pi) – same as score 2.5; weight loss 11- 20%; Score 4 (pi) - animal has ruffled fur or is positioned on its side or back, dehydrated, has difficulty breathing; weight loss >20%; Score 5 (pi) – death. On day four (Omicron variants BA.5 and XBB) or five (WT) post-challenge, all animals were euthanized and subjected to necropsy to remove the lungs. Left lungs were sectioned for hematoxylin and eosin (H&E) staining. Right lungs were homogenized for measurement of viral titers. Vero E6 cells were infected by serial diluted lung homogenates and TCID_50_ titers were calculated by Reed-Munch method.

### *In vivo* treatment of B16-OVA melanoma

6-8 weeks old C57BL/6 female mice were inoculated with 0.5 × 10^6^ B16-OVA cells in the right flank. For early-stage tumors, OVA mRNA-loaded LNP or PLNP (5 µg/mouse, n = 4) were intramuscular injected on day 7 (tumor size∼ 30 mm^3^). Booster shots were injected on day 10 post tumor inoculation. LNP or PLNP treated groups were rechallenged with 0.5 × 10^6^ B16-OVA cells inoculation in the left flank on day 25. For advanced-stage tumors, OVA mRNA-loaded LNP or PLNP (5 µg/mouse, n = 9) were intramuscular injected on day 9 post tumor inoculation (tumor size ∼85 mm^3^). Booster shots were injected on day 11, day 13, and day 15 post tumor inoculation. Tumor size and mouse survival were monitored. Tumor sizes were calculated using the following formula: height × width × width /2. The mice were euthanized when they reach the endpoint (tumor size reaches 1000 mm^3^).

## ACKNOWLEDGMENTS

This work was supported by NIH 3DP2AI144245-01S1, Walder Foundation, and Lynn Sage Research Award (to J.H.). We thank the University of Chicago Advanced Electron Microscopy Facility for support with TEM imaging and Soft Matter Characterization Facility for support with DLS. We thank the University of Chicago Integrated Small Animal Imaging Research Resource for support with bioluminescence and fluorescence imaging. We acknowledge the NIH Tetramer Core Facility for providing MHC monomers and tetramers. We thank the University of Chicago Flow Cytometry Core, Human Disease and Immune Discovery Core, and Dr. Cezary Ciszewski for expert support with flow cytometry experiments. We thank Pritzker School of Molecular Engineering Single-Cell Immunophenotyping Core for assisting in the generation of single-cell sequencing data. We thank the Howard Taylor Ricketts Laboratory BSL-3 facility for supporting viral neutralization and challenging experiments. We thank Kevin Chang, Dr. Jeffrey A. Hubbell, and the Hubbell lab for providing the B16-OVA cells and supporting this project. We further acknowledge the Animal Resources Center for assistance with animal studies and the University of Chicago Human Tissue Resource Center for support with histopathological analysis of tissues.

## AUTHOR CONTRIBUTION

M. C., X. C., and J. H. conceived the ideas and designed the project. J. H. supervised the project. X. C. and M. C. designed and synthesized the nanoparticles and conducted the *in vitro* characterization of nanoparticles. T. D. and P. P.-M. generated and provided the SARS-CoV-2 spike plasmid DNA vectors for mRNA vaccines and the spike protein used as coating antigen for ELISA, and provided advisory input on nanoparticle production, mouse vaccinations, and ELISA experiments. X. C., M. C., and N. A. designed and cloned plasmids for the mRNA constructs. T. P. assisted with mRNA sequence design and optimization. X. C., M. C. and R. S. synthesized the mRNAs. Q. C., Z. G. and A.E.K. provided BMDCs. X. C and D.-T. N. performed *in vivo* characterization of mRNA translation. X. C., M. C., X. H., N. A., W. Z., E. T., S. W., A. S., M. N., L. V., and R.W. performed *in vitro* and *in vivo* characterization of T cell and B cell responses post vaccination. X. C., M. C., and X. H. performed single-cell RNA sequencing experiments. G. C. analyzed the single-cell RNA sequencing data. X. C., M. C., A. S., H. G., D. E., V. N. performed the SARS-CoV-2 viral neutralization and viral challenge experiments. D. M. coordinated and provided advisory input on the SARS-CoV-2 viral neutralization and viral challenge experiments. X. C. and S. W. performed the *in vivo* B16-OVA tumor treatment experiments. X. C., M. C., G. C., H. G., and D. E. analyzed the data. X. C., M. C., and J. H. wrote the manuscript. J. H. acquired the funding.

