## Supporting Information for "Polymer-lipid hybrid nanoparticle enhances mRNA delivery and T cell-mediated immunity"

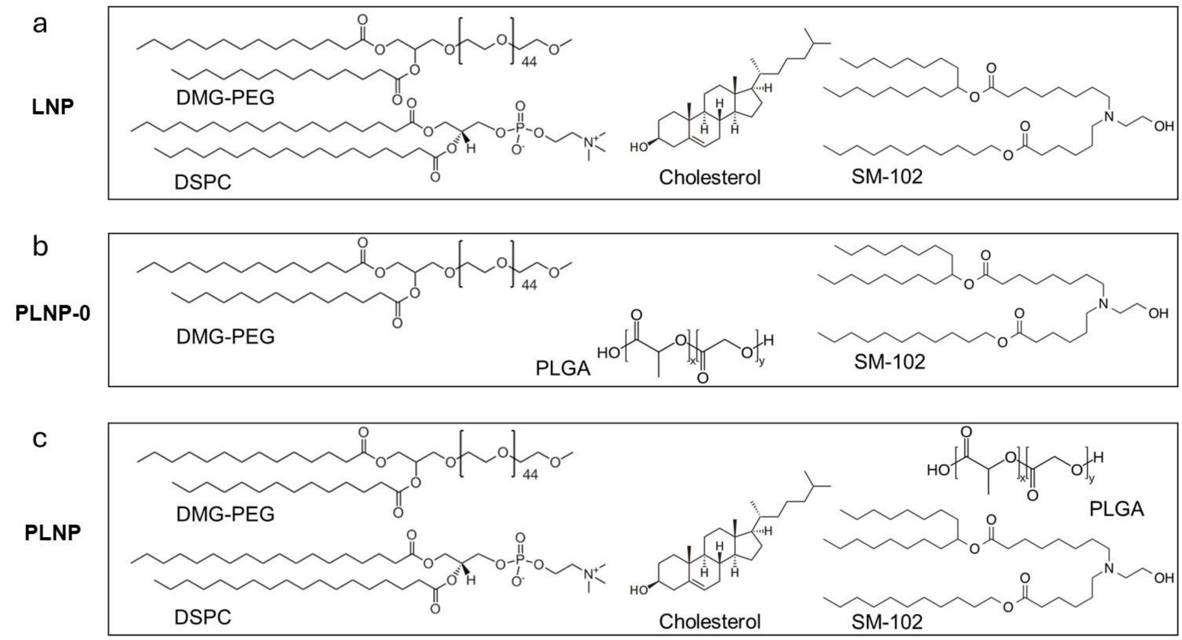


**Scheme S1**. **Components of LNP, PLNP-0, and PLNP.** (a) Components of LNP. (b) Components of PLNP-0. (c) Components of PLNP.

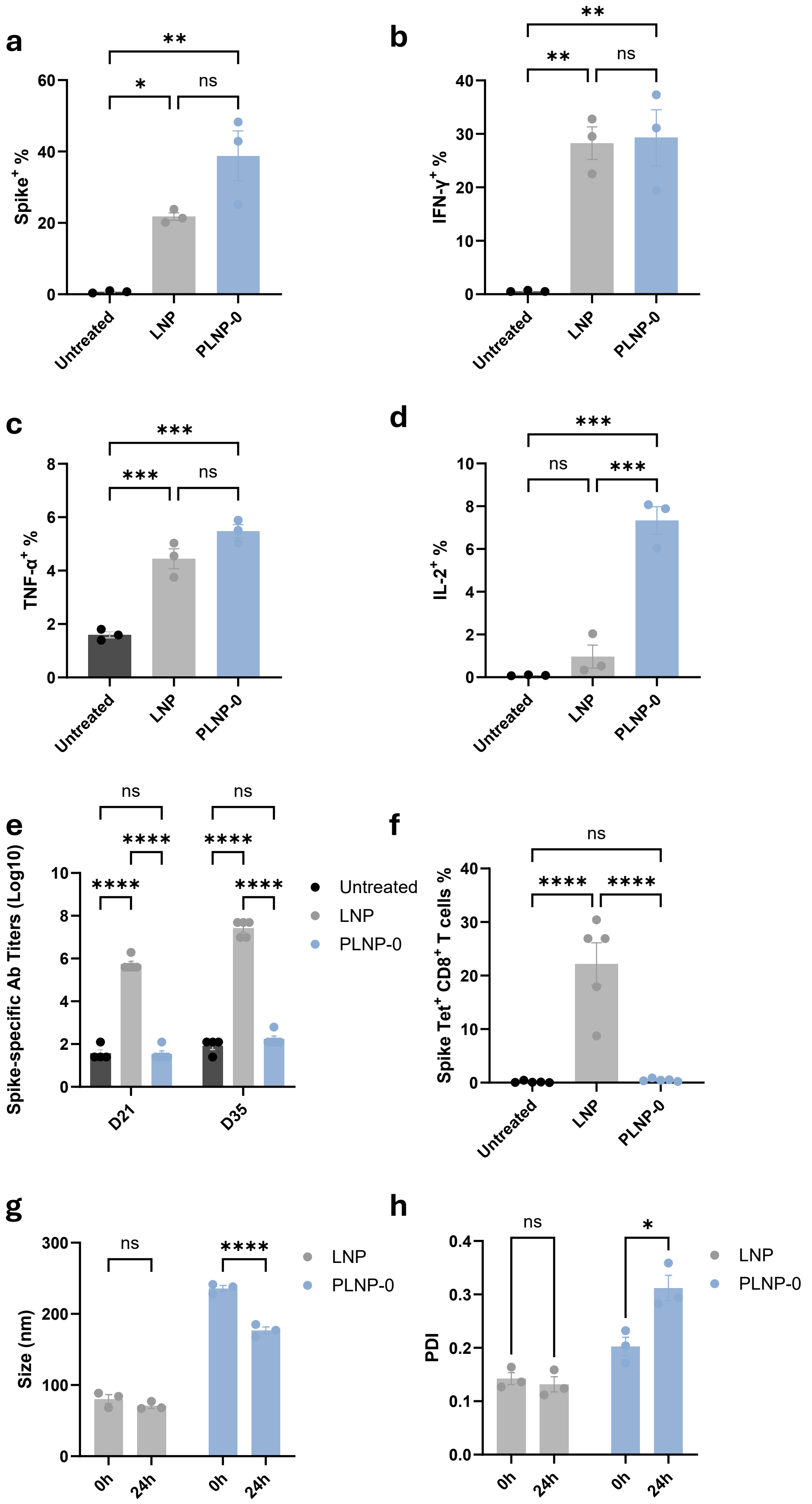
a
Spike+ %
Untreated
LNP
PLNP-0
**
*
ns
b
IFN-γ+ %
Untreated
LNP
PLNP-0
**
**
ns
c
TNF-α+ %
Untreated
LNP
PLNP-0
***
***
ns
d
IL-2+ %
Untreated
LNP
PLNP-0
***
ns
***
e
Spike-specific Ab Titers (Log10)
D21
D35
ns
ns
****
****
****
****
Untreated
LNP
PLNP-0
f
Spike Tet+ CD8+ T cells %
Untreated
LNP
PLNP-0
ns
****
****
g
Size (nm)
0h
24h
0h
24h
ns
****
LNP
PLNP-0
h
PDI
0h
24h
0h
24h
ns
*
LNP
PLNP-0

**Figure S1.** ***In vitro*** **characterization of LNP and PLNP-0.** (a) Spike mRNA transfection in RAW 264.7 macrophages after incubation with spike mRNA-loaded LNP and PLNP-0. (b-d) Intracellular cytokine levels in RAW 264.7 macrophages after incubation with spike mRNA -loaded LNP and PLNP-0 measured by intracellular staining and flow cytometry. (e) Anti-spike antibody generation on day 21 and day 35 after vaccinating C57BL/6 mice with spike mRNA-loaded LNP and PLNP-0. (f) ) The percentages of spike antigen-specific T cells in total $CD8^+$ T cells on day 28 after vaccinating C57BL/6 mice with spike mRNA-loaded LNP and PLNP-0. (g-h) Size and PDI changes after storing LNP and PLNP-0 at 4°C for 24 hours. Data are presented as mean ± s.e.m. Statistical significance was determined using one-way ANOVA with Tukey post hoc test for multiple comparisons (a, b, c, d, f), or two-way ANOVA with Tukey post hoc test for multiple comparisons (e, g, h). * $P < 0.05$, ** $P < 0.01$, *** $P < 0.001$, and ****$P < 0.0001$. n = 3.

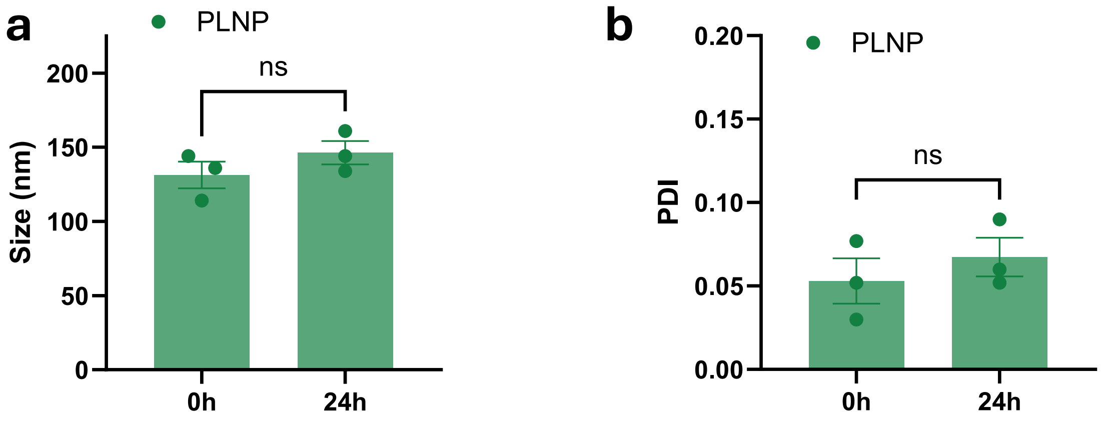


**Figure S2. In vitro characterization of PLNP.** (a-b) Size and PDI changes after storing PLNP at 4°C for 24 hours. Data are presented as mean ± s.e.m. Statistical significance was determined using two-sided unpaired t-test. ns: no significance. n = 3.

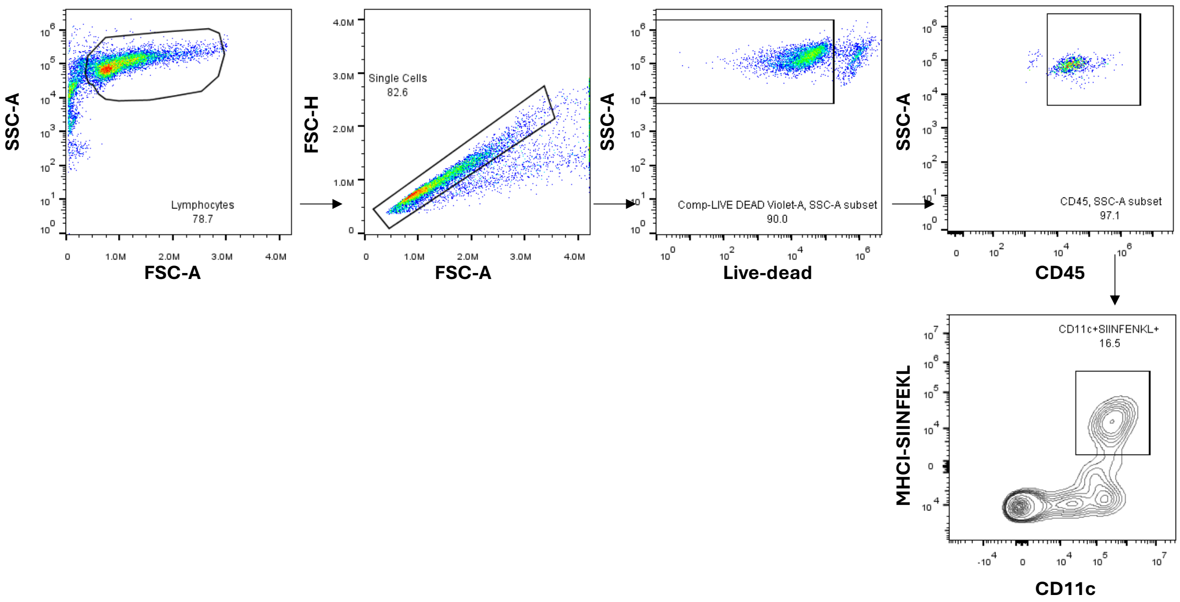


**Figure S3. Flow cytometry gating strategy for quantifying SIINFEKL/H-2K$^{b}$-positive DCs.** C57BL/6 mouse bone marrow-derived dendritic cells were incubated with OVA mRNA-loaded LNP or PLNP for 48 hours. The antigen-presenting DCs were quantified by gating CD45$^{+}$CD11c$^{+}$ SIINFEKL/H-2K$^{b+}$ DCs using flow cytometry. Dose: 1 μg mRNA/mL. n = 3.

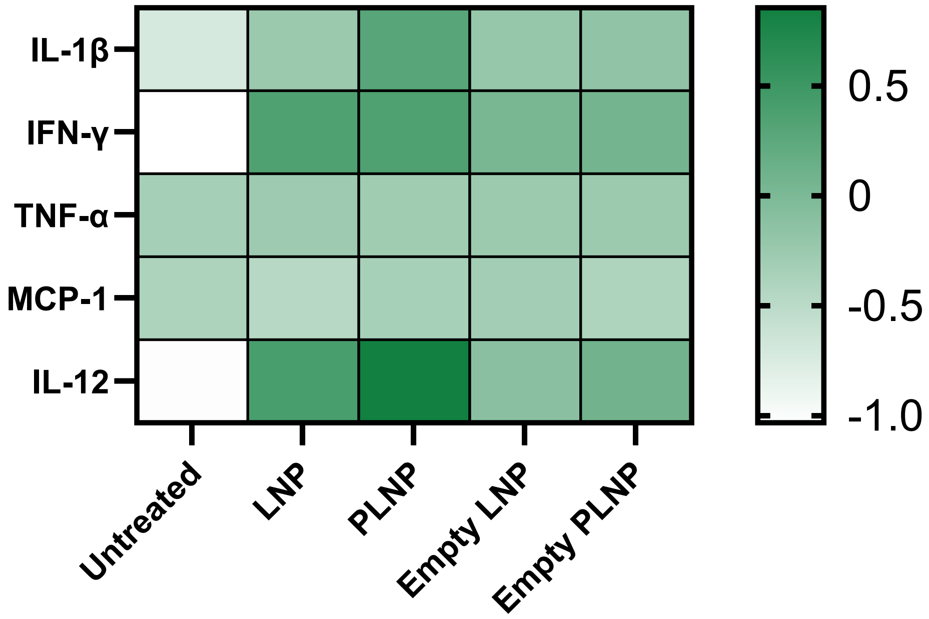


**Figure S4.** ***In vitro*** **proinflammatory cytokine secretion from mouse PBMCs after incubation with different formulations.** Mouse PBMCs were incubated with spike mRNA-loaded LNP, spike mRNA-loaded PLNP, empty LNP, or empty PLNP for 24 hours and cytokine secretion was measured by Legendplex assay. Dose: 1 µg mRNA/mL. n = 5.

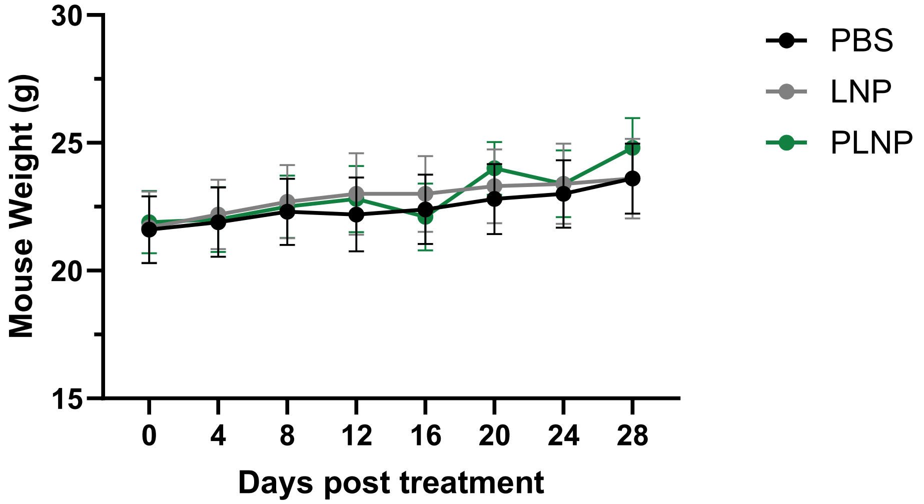


**Figure S5. Weight changes of C57BL/6 mice after LNP or PLNP treatment**. Mice were vaccinated with spike mRNA-loaded LNP or PLNP on day 0 and day 21 (10 µg mRNA/mouse) and their body weight were measured from day 0 to day 28. Data are presented as mean ± s.e.m. n = 10.

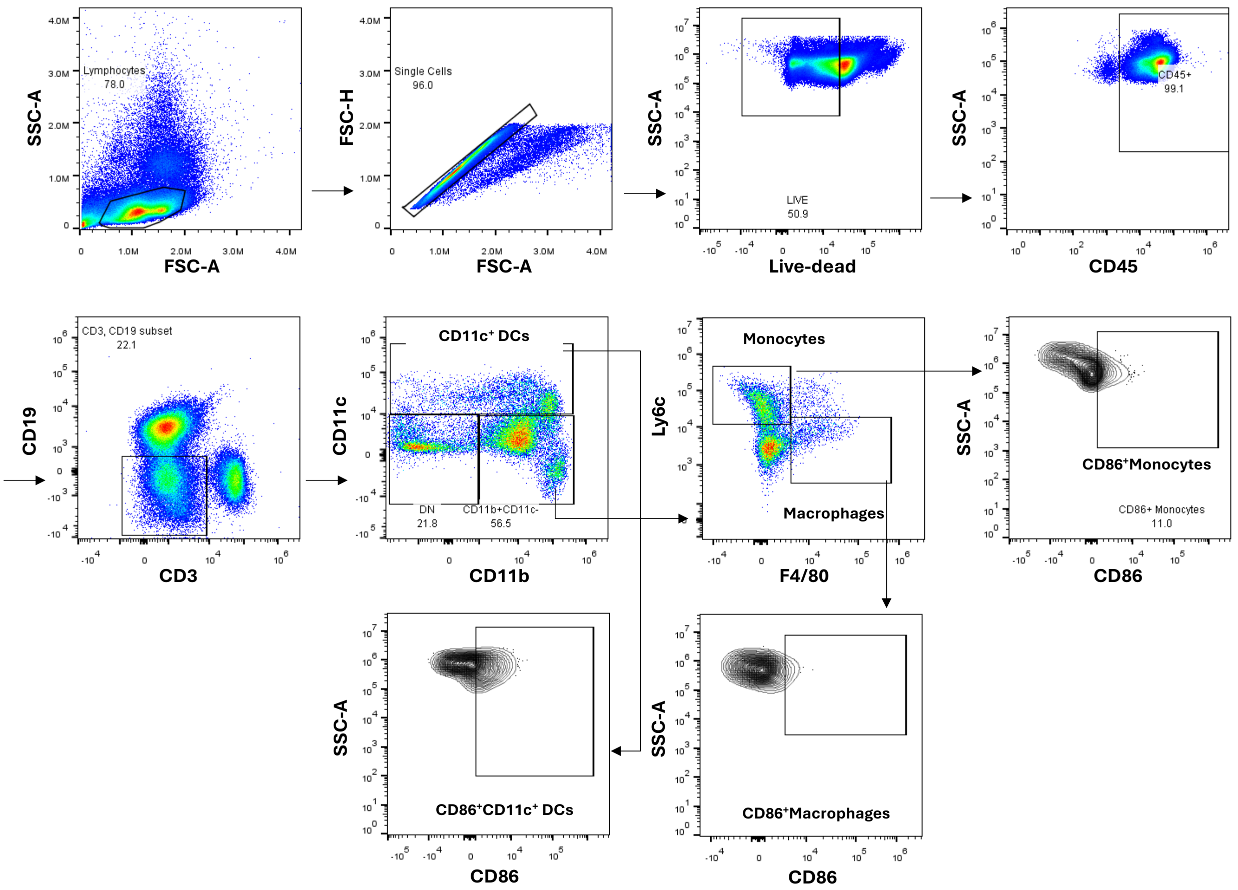


**Figure S6. Flow cytometry gating strategy for characterizing innate immune cells in draining lymph nodes.** C57BL/6 mice were vaccinated with spike mRNA-loaded LNP or PLNP on day 0 and day 21. Dose: 10 µg mRNA/mouse. n = 5. Draining lymph nodes were collected on day 28 for innate immune cell characterization by gating DCs ($CD45^+CD3^-CD19^-CD11c^+$), macrophages ($CD45^+CD3^-CD19^-CD11c^-CD11b^+Ly6c^-F4/80^+$), and monocytes ($CD45^+CD3^-CD19^-CD11c^-CD11b^+Ly6c^+F4/80^-$) using flow cytometry.

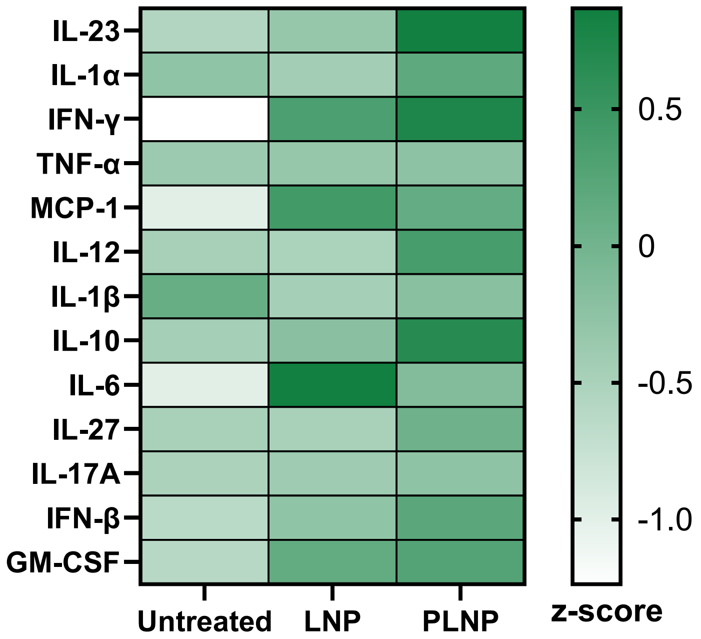


**Figure S7. In vivo systematic proinflammatory cytokine levels induced by LNP or PLNP**. C57BL/6 mice were vaccinated with spike mRNA-loaded LNP or PLNP. Dose: 10 μg mRNA/mouse. n = 5. Serum samples were collected at 24 h post vaccination for cytokine level measurement by Legendplex assay.

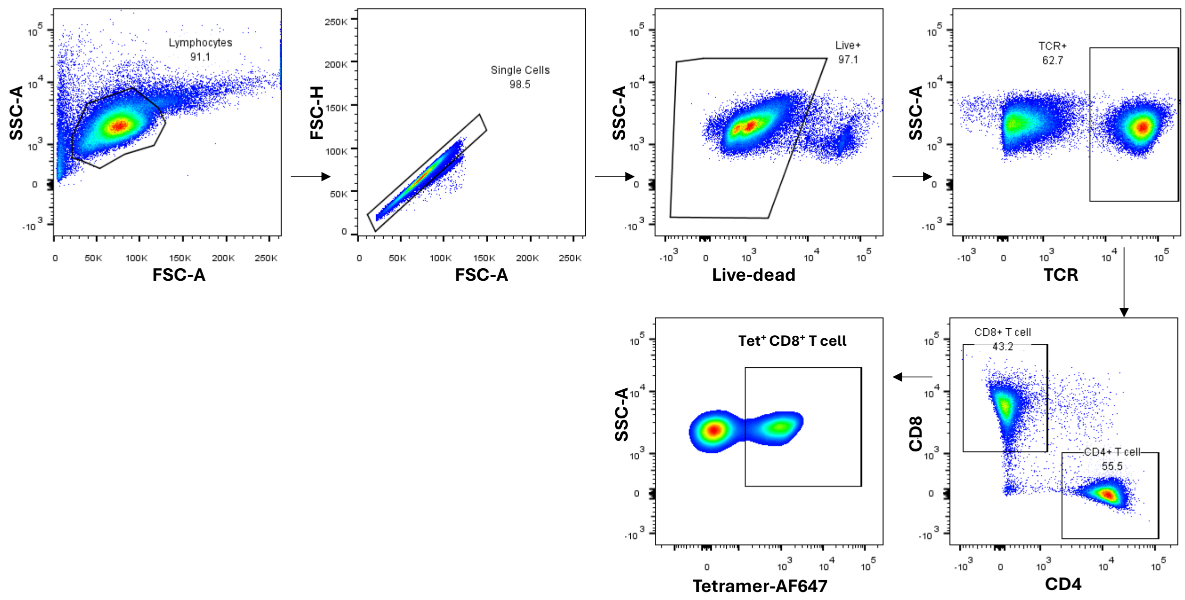


**Figure S8. Flow cytometry gating strategy for characterizing T cells in spleens.** C57BL/6 or BALB/c mice were vaccinated with spike, HA, or OVA mRNA-loaded LNP or PLNP on day 0 and day 21. Dose: 10 μg spike mRNA, 10 μg HA mRNA, or 5 μg OVA mRNA/mouse. n = 5. Splenocytes were collected on day 28 for T cell characterization by gating $TCR^{+}CD8^{+}Tetramer^{+}$ T cells.

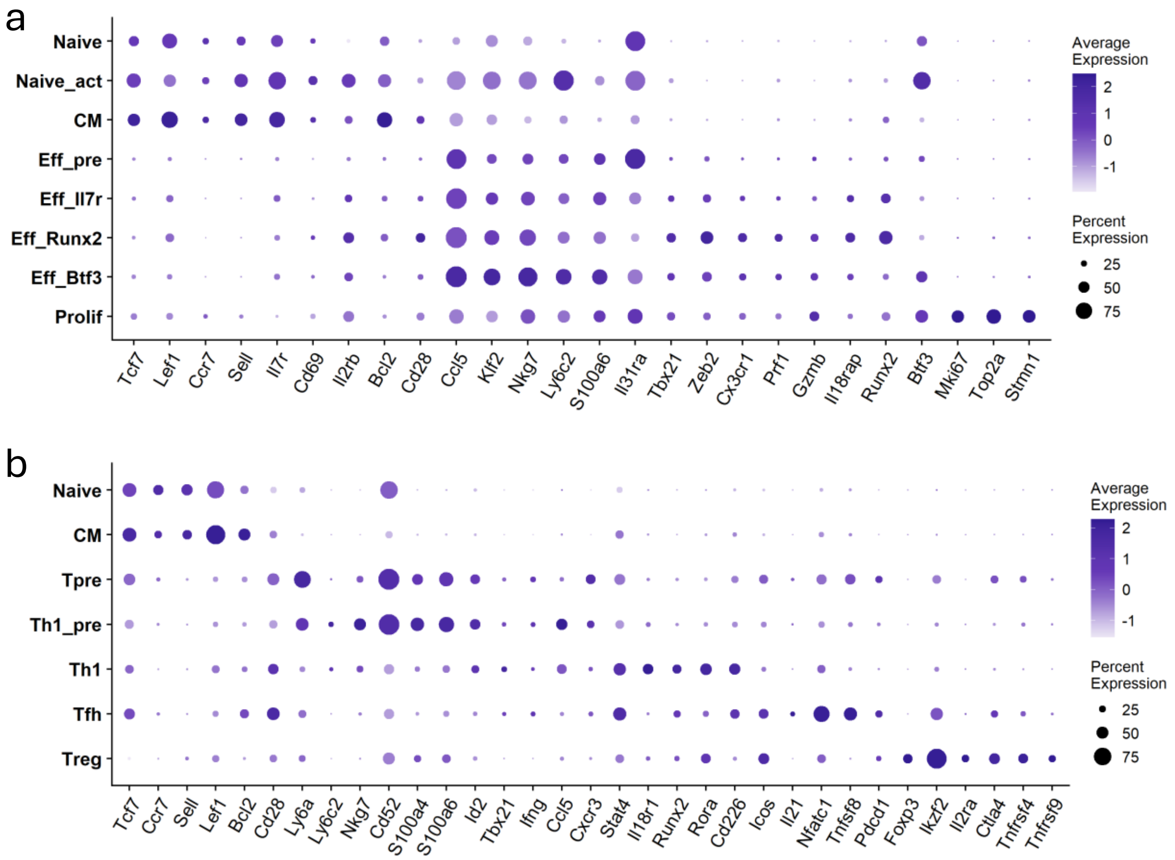


**Figure S9. T-cell characterization by scRNA-seq post vaccination with LNP and PLNP.** C57BL/6 mice were vaccinated with spike mRNA-loaded LNP or PLNP on day 0 and day 21. Dose: 10 µg mRNA/mouse. n = 5. Splenocytes were collected on day 28 for T cell characterization. **(a)** Dot plot of cluster-specific markers of $CD8^+$ T cells specifically upregulated in each cluster compared to other clusters. These markers were predominantly used to define the phenotypes of each cluster in $CD8^+$ T cells. **(b)** Dot plot of cluster-specific markers of $CD4^+$ T cells specifically upregulated in each cluster compared to other clusters in $CD4^+$ T cells. These markers were predominantly used to define the phenotypes of each cluster in $CD4^+$ T cells.

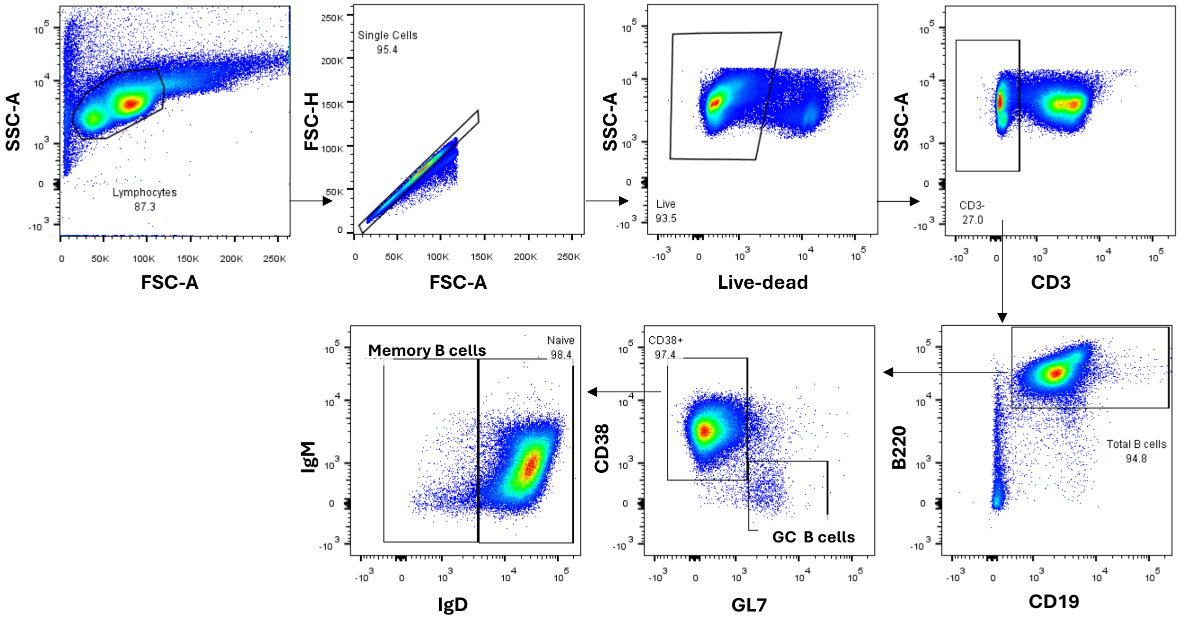


**Figure S10. Flow cytometry gating strategy for characterizing B cells in draining lymph nodes.** C57BL/6 mice were vaccinated with spike mRNA-loaded LNP or PLNP on day 0 and day 21. Dose: 10 μg mRNA/mouse. n = 5. Draining lymph nodes were collected on day 28 for B cell characterization by gating GC B cells ($CD3^{-}CD19^{+}B220^{+}CD38^{-}GL7^{+}$) and memory B cells ($CD3^{-}CD19^{+}B220^{+}CD38^{+}GL7^{-}IgD^{-}$) using flow cytometry.

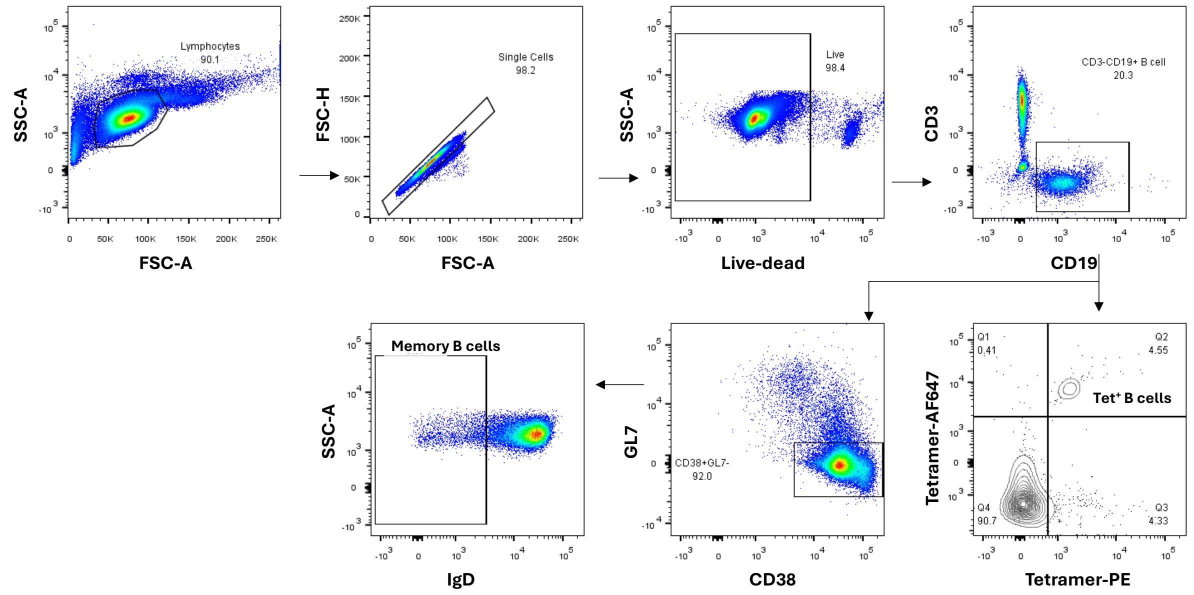


**Figure S11. Flow cytometry gating strategy for characterizing B cells in spleens.** C57BL/6 mice were vaccinated with spike mRNA-loaded LNP or PLNP on day 0 and day 21. Dose: 10 µg mRNA/mouse. n = 5. Splenocytes were collected on day 28 for B cell characterization by gating memory B cells ($CD3^-CD19^+CD38^+GL7^-IgD^-$) and antigen-specific B cells ($CD3^-CD19^+$Tetramer-$PE^+$Tetramer-$AF647^+$) using flow cytometry.

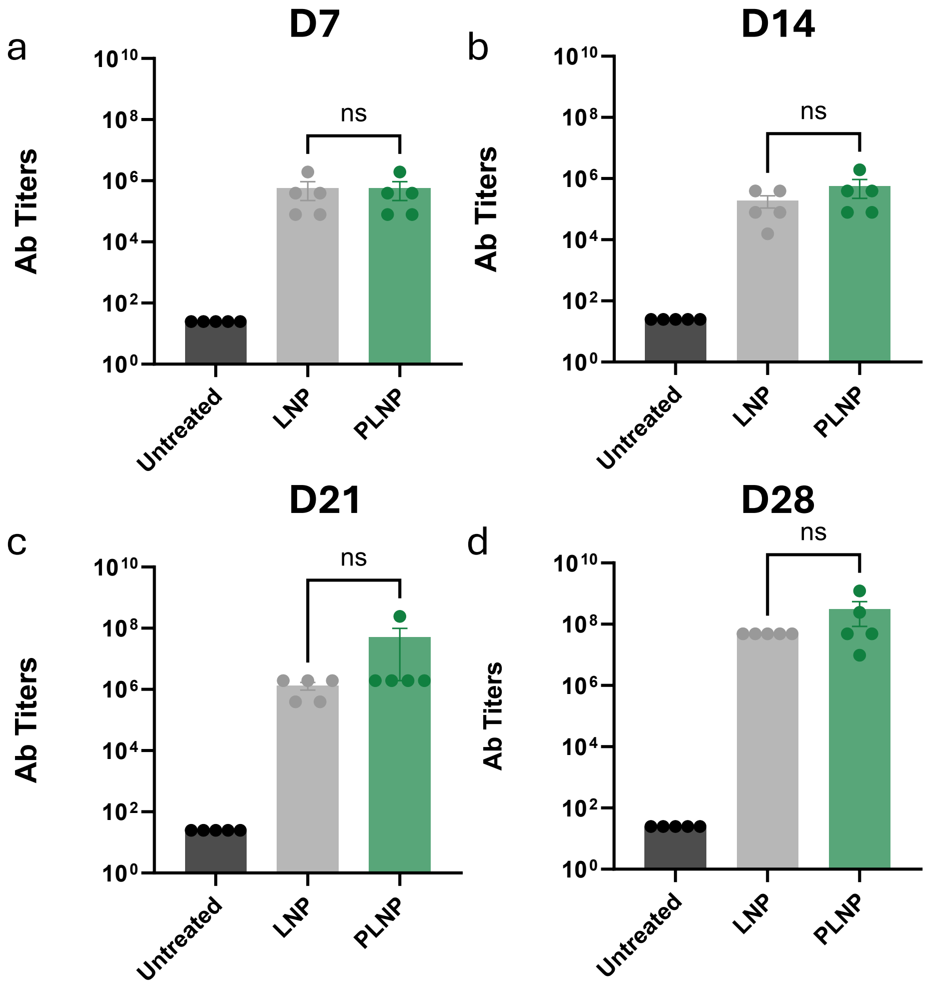


**Figure S12. HA-specific antibody production post vaccination with influenza HA mRNA-loaded LNP or PLNP.** BALB/c mice were vaccinated with influenza HA mRNA loaded LNP or PLNP on day 0 and day 21. Dose: 10 µg mRNA/mouse. n = 5. Serum samples were collected on day 7 (a), day 14 (b), day 21 (c) and day 28 (d) for antibody quantification by ELISA. Data are presented as mean ± s.e.m. Statistical significance was determined using two-sided unpaired t-test. ns: no significance.

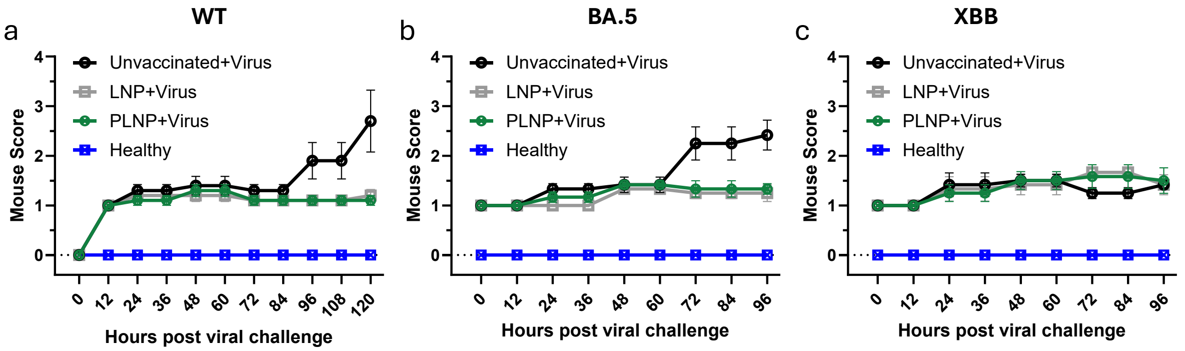


**Figure S13. Mouse clinical scores post SARS-CoV-2 viral challenge.** Mice were vaccinated with spike mRNA-loaded LNP or PLNP on day 0 and day 21 and further challenged with different SARS-CoV-2 variants (WT, BA.5 and XBB) on day 35. Dose: 10 µg mRNA/mouse. n = 5. Mice were monitored twice a day and clinical scores were recorded. Score 0 (pre-inoculation) - animal is bright, alert, active, with normal fur coat and posture; Score 1 (post-inoculation, pi) – animal is bright, alert, active, normal fur coat and posture, no weight loss; Score 1.5 - animal has slightly ruffled fur but is active; weight loss under 2.5%; Score 2 (pi) – animal has ruffled fur, is less active; weight loss under 5%; Score 2.5 (pi) - animal has ruffled fur, is not active but moves when touched, may have hunched posture or difficulty breathing; weight loss 5-10%; Score 3 (pi) – same as score 2.5; weight loss 11- 20%; Score 4 (pi) - animal has ruffled fur or is positioned on its side or back, dehydrated, has difficulty breathing; weight loss >20%;  Score 5 (pi) – death.
